# Nicotine exposure and neuronal activity regulate Golgi membrane dispersal and distribution

**DOI:** 10.1101/2020.02.25.965285

**Authors:** Anitha P. Govind, Okunola Jeyifous, Theron A. Russell, Lee O. Vaasjo, Zola Yi, Aubrey V. Weigel, Luke Newell, Jessica L. Koranda, Karanveer Singh, Fernando Valbuena, Benjamin S. Glick, Jogeshwar Mukherjee, Jennifer Lippincott-Schwartz, Xiaoxi Zhuang, William N. Green

**Affiliations:** Department of Neurobiology, University of Chicago, Chicago IL, 60637; Department of Molecular Genetics and Cell Biology, University of Chicago, Chicago, IL 60637; Preclinical Imaging, Department of Radiological Sciences, University of California-Irvine, Irvine, CA 92697; Janelia Research Campus, Howard Hughes Medical Institute, Ashburn, Virginia 20147; Marine Biological Laboratory, Woods Hole, MA 02543

## Abstract

How nicotine exposure produces long-lasting changes that remodel neural circuits with addiction is unknown. Here, we report that long-term nicotine exposure alters the trafficking of α4β2-type nicotinic acetylcholine receptors (α4β2Rs) by dispersing and redistributing the Golgi apparatus. In cultured neurons, dispersed Golgi membranes were distributed throughout somata, dendrites and axons. Small, mobile vesicles in dendrites and axons lacked standard Golgi markers and were identified by other Golgi enzymes that modify glycans. Nicotine exposure increased levels of dispersed Golgi membranes, which required α4β2R expression. Similar nicotine-induced changes occurred *in vivo* at dopaminergic neurons at mouse nucleus accumbens terminals, consistent with these events contributing to nicotine’s addictive effects. Characterization *in vitro* demonstrated that dispersal was reversible, that dispersed Golgi membranes were functional, and that membranes were heterogenous in size, with smaller vesicles emerging from larger “ministacks”, similar to Golgi dispersal induced by nocadazole. Protocols that increased cultured neuronal synaptic excitability also increased Golgi dispersal, without the requirement of α4β2R expression. Our findings reveal novel activity- and nicotine-dependent changes in neuronal intracellular morphology. These changes regulate levels and location of dispersed Golgi membranes at dendrites and axons, which function in local trafficking at subdomains.

## Introduction

Tobacco use continues world-wide and is the leading cause of preventable deaths in the United States (U.S. Department of Health and Human Services 2014). Nicotine, the addictive molecule in tobacco, binds to high-affinity nicotinic acetylcholine receptors (nAChRs) in brain, where it initiates its addictive effects. nAChRs are members of the Cys-loop family of ligand-gated ion channels, all of which are pentameric neurotransmitter receptors (Karlin 2002, Albuquerque, Pereira et al. 2009). In mammalian brains, high-affinity nicotine binding sites are largely nAChRs containing α4 and β2 subunits (Albuquerque, Pereira et al. 2009). In mice, knockout of either subunit reduced the pharmacological and behavioral effects of nicotine (Picciotto, Zoli et al. 1998, Marubio, Gardier et al. 2003), which can be rescued by targeted β2 subunit expression in the mid-brain reward ventral tegmental area (VTA; (Maskos, Molles et al. 2005). Other studies link α4β2 nAChRs (α4β2Rs) to nicotine addiction (Vezina, McGehee et al. 2007, Govind, Vezina et al. 2009, Govind, Walsh et al. 2012, Lewis and Picciotto 2013), which involves the process of nicotine upregulation of α4β2Rs. Chronic nicotine exposure causes α4β2R upregulation, a complex set of long-term changes causing both increases in α4β2R high-affinity binding site numbers and functional upregulation, the increased α4β2R functional response after nicotine exposure (Marks, Burch et al. 1983, Schwartz and Kellar 1983, Benwell, Balfour et al. 1988, Breese, Adams et al. 1997).

Recently, we described how the anti-smoking drug, varenicline (Chantix), is selectively trapped as a weak base within intracellular acidic vesicles that contain high-affinity α4β2Rs (Govind, Vallejo et al. 2017). Trapping maintains levels of varenicline throughout the day in the brain and appears to act when nicotine levels decline. Varenicline trapping is amplified by two processes involving nicotine upregulation. First, nicotine exposure increases the number of high-affinity binding sites, which increases the capacity of the acidic vesicles to trap varenicline. Second, nicotine exposure increases the number of the acidic vesicles further amplifying the effects. In this study, we address the nature of these α4β2R-containing acidic vesicles, where varenicline is trapped, and how these vesicles arise.

Here, we find that α4β2R-containing acidic vesicles result from dispersal of the Golgi apparatus. The morphology and positioning of the Golgi changes during neuronal development and has critical a role in establishing neuronal polarity (Zmuda and Rivas 1998, Horton, Racz et al. 2005, de Anda, Meletis et al. 2010). What happens to the Golgi after neuronal development is less well characterized. mRNAs for secreted and membrane proteins, along with cytoplasmic protein mRNA, are transported to sites at specialized domains in dendrites and axons where localized protein translation appears to occur. However, the Golgi elements that usually mediate the last steps in the processing of secreted and membrane proteins have not been observed in axons and appear to be largely absent in dendrites (Hanus and Ehlers 2016). This lack of Golgi in dendrites has led to the idea that unconventional secretory pathways or “Golgi bypass” replaces Golgi membranes at sites of local translation in dendrites (Hanus and Ehlers 2008). Evidence supportive of an unconventional secretory pathway at sites of local translation in dendrites is the finding of high levels of high-mannose or “immature” N-linked glycans on the surface of cultured neurons (Hanus and Ehlers 2016, Bowen, Bourke et al. 2017). A different study suggests that an unconventional secretory pathway in dendrites may traffic through recycling endosomes instead of Golgi membranes (Bowen, Bourke et al. 2017)

Contrary to findings of a lack of dendritic Golgi membranes, others have observed Golgi membranes in addition to Golgi “outposts” in dendrites (Mikhaylova, Bera et al. 2016) (Stoeber, Jullie et al. 2018). The trafficking of newly synthesized cargo from ER to dendritic Golgi membranes has been observed for NMDA-type glutamate receptors (Jeyifous, Waites et al. 2009), alpha7-type nAChRs (Alexander, Sagher et al. 2010) and GluK2-containing kainate receptors (Evans, Gurung et al. 2017). In support of these studies, here we find that dispersed Golgi membranes are much more widely distributed in neurons than previously characterized. The dispersed elements were not widely observed previously because the standard Golgi markers originally used are not present in smaller dispersed Golgi membranes. We observed dispersed Golgi membranes in the somata, dendrites and axons of neurons. A subset are smaller dispersed Golgi vesicles that are highly mobile, moving at a rate of ∼1 μm/sec. Nicotine exposure increases Golgi dispersal throughout cultured neurons and in ventral tegmental area (VTA) dopaminergic neuron terminals in the mouse nucleus accumbens, thereby increasing the number of dispersed Golgi membranes ∼2-fold in dendrites and axons. Nicotine-induced Golgi dispersal does not appear to have the pathological effects linked to different neurodegenerative diseases, as it is reversible, and the dispersed elements are functional. Increases in synaptic activity also increase Golgi membranes ∼2-fold in dendrites and axons. However, the effects of nicotine only occur with α4β2R expression, while the effects of synaptic activity occur in their absence. Our findings reveal novel long-lasting changes in GA morphology and the distribution of Golgi vesicles throughout neurons driven by nicotine exposure and synaptic activity.

## Results

### Golgi dispersal by nicotine in cells stably expressing α4β2Rs

In this study, our goal was to identify the intracellular acidic vesicles that contain high-affinity α4β2Rs and trap varenicline (Govind, Vallejo et al. 2017). We examined the Golgi apparatus (GA), which gives rise to an acidic compartment in cells (Casey, Grinstein et al. 2010). In HEK cells stably expressing α4β2Rs, we observed that nicotine exposure for 17-18 hours altered the morphology of the Golgi apparatus by changing the perinuclear compact stacked structure into a dispersed set of membranes throughout the cytoplasm (Fig. 1A). These changes were initiated at nicotine concentrations of 300 nM or below and saturated at ∼1 μM nicotine, similar to the nicotine dose-dependence of nicotine upregulation in these cells (Vallejo, Buisson et al. 2005, Govind, Walsh et al. 2012). We observed no changes in Golgi morphology without α4β2R expression in the HEK cells (Fig. 1B), and thus conclude that α4β2R expression is required for the nicotine-dependent dispersal to occur.

**Figure 1.**
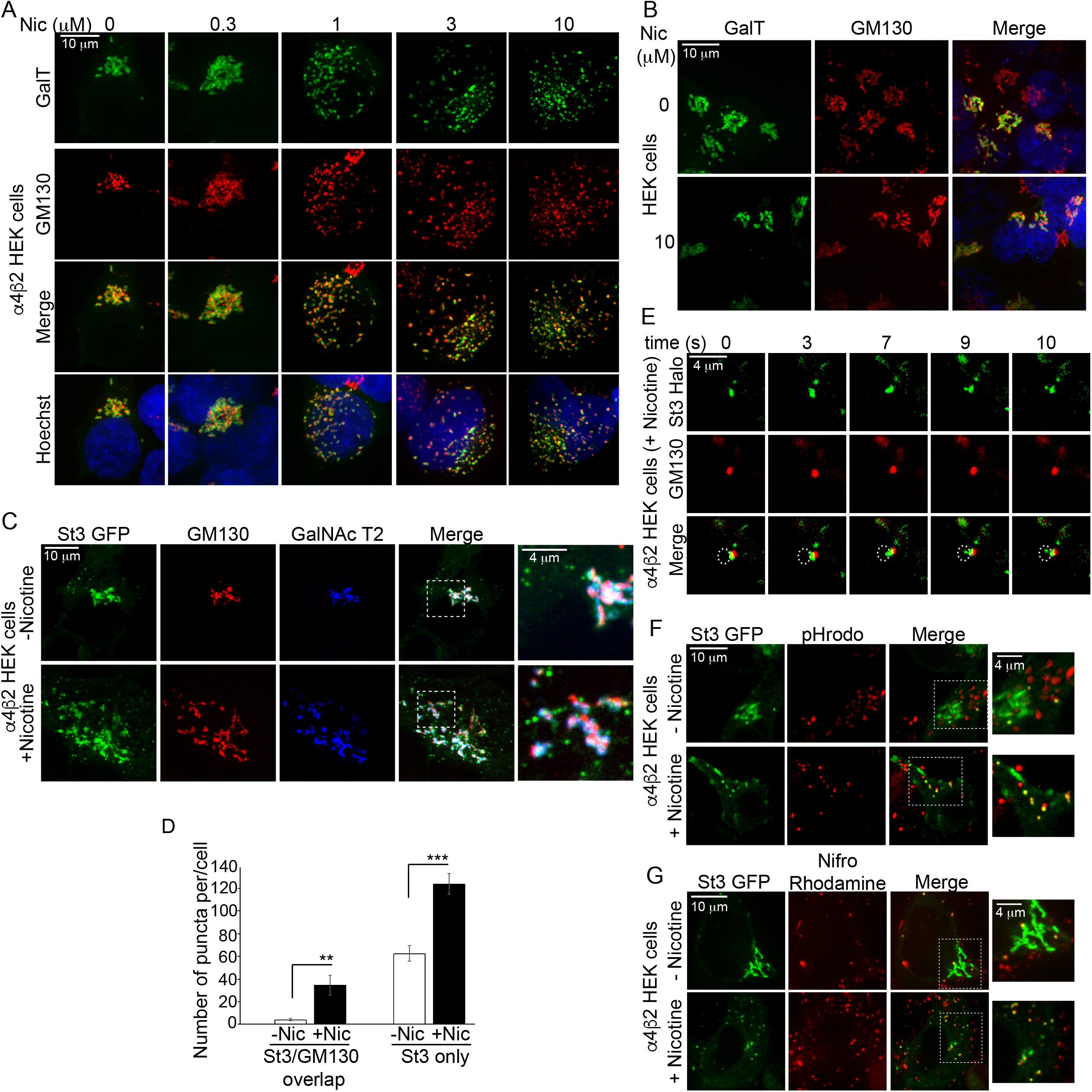
A. Dose dependence of nicotine-induced Golgi dispersal. HEK cells that stably express α4β2 receptors (α4β2 HEK cells) were transfected with triple GFP-tagged galactosyl transferase (GalT, green) for 24 hours, treated with indicated concentrations of nicotine for 17 hours, and labeled with anti-GM130 antibody (red; secondary antibody anti-mouse Alexa 568) and DAPI (Blue). Scale bar, 10 μm. B. Nicotine does not alter Golgi morphology in HEK cells that do not express α4β2 receptors. HEK cells lacking the nicotinic receptors were transfected with Galtase-GFP (GalT, green), treated with 10uM nicotine for 17 hours, and fixed and stained with anti-GM130 and DAPI as in A. Scale bar, 10 μm. C. Golgi resident proteins show varying degrees of dispersal by nicotine. α4β2 HEK cells were transfected with eGFP tagged N-acetylgalactoside α(2,3) sialyl transferase 3 (St3-GFP, green) and mCherry-tagged N-acetylgalactosaminyl transferase 2 (GalNAc T2, blue) and treated with 10 μM Nicotine for 17 hours prior to fixation and staining for GM130 (red). Scale bar, 10 μm. Inset scale bar, 4 μm. D. Quantification of puncta that possess both GM130 and St3, or St3 only. Total number of puncta per cell, data are shown as mean ± SEM. For left graph, control cells, 3.9 ± 1.1; nicotine cells, 34.6 ± 9.0 (n = 8-10 cells per group, **p<0.005). For right graph, control cells, 62.6 ± 6.9; nicotine cells, 124.1 ± 8.9 (n = 10 cells per group, ***p<0.00001). E. St3-only vesicles emerging from the trans face of polarized Golgi mini stacks. α4β2 HEK cells were transfected with St3-Halo (green) and GM130-GFP (red) were treated with 10 μM Nic for 17 hours and imaged live. Scale bar, 4 μm. F. Acidic nature of St3-containing vesicles. α4β2 HEK cells were transfected with St3-GFP (green) and treated with 10 μM nicotine for 17 hours. Cells were loaded with pH-sensitive pHrodo dye (red) for 20 minutes. Cells were washed and imaged live in the presence of 10 μM nicotine. Scale bar, 10 μm. Inset scale bar, 4 μm. G. α4β2 receptors concentrate in acidic St3-containing vesicles. α4β2 HEK cells were transfected with St3-GFP (green) and treated with 10 μM nicotine for 17 hours. Cells were incubated with a fluorescent nicotinic receptor ligand, 50 μM nifrorhodamine (red), for 30 minutes, then washed and imaged live. Scale bar, 10 μm. Inset scale bar, 4 μm.

The features of the Golgi dispersal by nicotine varied depending on the Golgi “marker” used to label Golgi membranes. The observed differences in Golgi marker distributions were especially critical for experiments in neurons, where the Golgi in dendrites has generally been assayed using Abs specific for the cis-Golgi structural protein, GM130. We used a variety of different Golgi markers, including different cis-, medial- and trans-Golgi markers (Table I), to examine the differences in their distribution in HEK cells and neurons. Antibodies (Abs) specific for Golgi proteins at different locations in the polarized GA were used to assay endogenous proteins, and cDNA constructs of Golgi proteins for live imaging of “over-expressed” proteins. Fig. 1A shows Golgi morphology visualized with an Ab specific for GM130 and transfected trans-Golgi, GFP-tagged enzyme Beta-1,4,galactosyltransferase (GalT-GFP). Two types of dispersed elements were observed, larger membranes containing both markers and smaller membranes which only contained GalT-GFP. Additional images of the dispersed Golgi membranes are displayed in Fig. 1C and reveal more details of the heterogeneity of the membranes in terms of size and composition. Golgi membranes were imaged with other Golgi markers, including the GFP-tagged enzyme ST3 beta-galactoside alpha-2,3-sialyl transferase 3 (St3-GFP) and the mCherry-tagged enzyme N-acetylgalactosaminyl transferase 2 (GalNAcT2-mCherry), in addition to the cis-Golgi GM130 Ab. St3 and GalNAcT2 are trans- and medial-Golgi proteins, respectively (Roth, Taatjes et al. 1985) (Rottger, White et al. 1998). We observed all three Golgi proteins co-localizing within larger structures of the intact GA in the absence of nicotine treatment (top panels) and with nicotine treatment (lower panels). The three proteins did not perfectly align and were slightly offset because the intact GA and the larger nicotine-dispersed Golgi membranes are polarized. In addition to the larger structures, we observed smaller membrane structures containing only St3-GFP.

**Table 1.** List of antibodies (sources) and transfected cDNAs for the Golgi markers used in the current study. Sub-Golgi localization for each marker (cis, medial, or trans) is indicated.

| **Golgi markers** | **Sub-Golgi**<br>**Localization** | **Antibody** | **FP/epitope tag**<br>**(transfected)** |
| --- | --- | --- | --- |
| ST3 Beta-Galactoside<br>Alpha-2,3-Sialyltransferase 3<br>(St3) | Trans | Rb pAb (Abcam) | GFP, HaloTag |
| ST3 Beta-Galactoside<br>Alpha-2,3-Sialyltransferase 5<br>(St3Gal5) | Medial, Trans | Rb pAb, (Tiemeyer lab) | N/A |
| Golgi Matrix Protein (GM130) | Cis | Ms mAb (BD transduction);<br>Rb pAb (Sigma) | GFP |
| Beta-1,4-galactosyltransferase<br>(GalT) | Trans | N/A | 3xGFP |
| Polypeptide<br>N-Acetylgalactosaminyltransferase 2<br>(GalNAc T2) | Medial | N/A | GFP, mCherry |
| Mannosidase Alpha Class 2A Member 1<br>(Man) | Medial | N/A | GFP |

Finer details of the St3-GFP structures were captured using structured illumination microscopy (SIM; supplemental Fig. S1-1A). We analyzed the number and size distribution of the Golgi membranes labeled with St3-GFP and GM130 Ab in nicotine-treated and untreated cells (Fig. 1D). Membranes were classified as either both St3- and GM130-containing (St3/GM130 overlap) or St3 alone (St3 only). As shown in the size distribution histogram (Supplemental Fig. S1-1B), the sizes of the two different types of Golgi membranes were heterogenous, but St3-only membranes were significantly smaller on average than the St3/GM130 membranes. St3-only membranes were also more abundant in terms of the number of puncta per cell (Fig. 1D, Supplemental Fig. S1-1B) and for both types of Golgi membranes this value was significantly increased by nicotine. Another difference between the two classifications of Golgi membranes was that St3/GM130 membranes were largely immobile while smaller St3 membranes were often mobile (Fig S1-2; video).

An example of an immobile, St3/GM130 Golgi membrane generated following nicotine treatment is shown in Fig. 1E. Here, Halo-tagged St3 (St3-Halo; red) and GFP-GM130 (green) were transfected to permit live-imaging of the Golgi membranes. St3-Halo and GFP-GM130 fluorescence overlapped on the same structure but with an offset (see merged bottom images), consistent with a cis/trans polarized Golgi structure, or “mini-stack”, as observed with Golgi dispersal after Nocodazole treatment (Fourriere, Divoux et al. 2016). In the time lapse shown in Fig. 1E, a mobile, smaller St3 membrane emerges from the trans end of the polarized St3/GM130 Golgi membrane and moves away from the mini-stacked Golgi membrane. We have observed several of these events, consistent with smaller St3 membranes largely originating from the trans end of the polarized Golgi structures labeled by St3 and GM130.

The degree of Golgi dispersal by nicotine varied between cells in HEK cell cultures. To quantify this variability, cells expressing St3-GFP in culture were binned based on whether the GA was intact, partially dispersed, or fully dispersed (Supplemental Fig. S1-1C), categories previously used to describe Golgi dispersal in cultured cells (Wortzel, Koifman et al. 2017). Examples of each category are displayed in Supplemental Fig. S1-1C. Without nicotine treatment, ∼70% of the HEK cells had a fully intact GA, ∼20% displayed a partially dispersed phenotype, and less than 10% had fully dispersed Golgi. Similar results were obtained previously with HeLa cells stained with an anti-GM130 Ab and binned by eye or by cell sorting (Wortzel, Koifman et al. 2017). For α4β2R-expressing HEK cells treated with nicotine, ∼20% of the cells had fully intact Golgi, ∼30% exhibited partial dispersal and ∼50% had fully dispersed Golgi. Dispersal variability appears to be largely caused by cell-to-cell differences in α4β2R expression across the cultures, a feature that increases with cell passage number, probably because the cell cycle is slowed by α4β2R expression. Golgi dispersal during the cell cycle may also be a contributing factor.

To assess whether St3-containing dispersed Golgi membranes and α4β2R-containing acidic vesicles (Govind et al. 2017) constitute a shared compartment, we tested to see if Golgi membranes are acidic and contain high-affinity α4β2Rs. We co-stained with pHrodo, a fluorescent probe that stains membranous compartments which have a lumenal pH lower than 6. To image high-affinity α4β2Rs, we developed a membrane-permeant version of a fluorescent high-affinity probe that binds α4β2Rs, nifrorhodamine. Nifrorhodamine’s structure is based on a different membrane-impermeant probe, nifrofam (Samra, Intskirveli et al. 2018) and specifically labels high-affinity α4β2Rs, which are only a small subset of the total α4 and β2 subunits and concentrate in acidic vesicles(Govind, Vallejo et al. 2017).

Fig. 1F shows images of α4β2R-expressing HEK cells transfected with St3-GFP (green) and stained with pHrodo (red). In the absence of nicotine treatment (top), only a few of the St3-containing puncta are co-stained by pHrodo, and the number of co-stained puncta was increased in cells treated with nicotine (bottom). We observed a similar staining pattern with nifrorhodamine, namely an increase in co-staining of St3-containing puncta with nicotine treatment (Fig. 1G). Both pHrodo and nifrorhodamine only stained a subset of the small St3-containing puncta. This finding was consistent with the numbers of St3-containing puncta per cell (Fig. 1D) being greater than the number of α4β2R-containing acidic puncta per cell previously measured (Govind et al. 2017). Based on the staining profile of St3-containing puncta with pHrodo and nifrorhodamine, we conclude that St3-containing dispersed Golgi membranes do share features of the α4β2R-containing acidic vesicles.

### Golgi dispersal by nicotine in cultured neurons expressing α4β2Rs

Next, we examined Golgi membranes in cultured neurons from rat cortex or hippocampus in which α4β2Rs were overexpressed. Only 5-10% of cultured cortical neurons endogenously express α4β2Rs, and their expression is largely observed in inhibitory neurons (Govind, Walsh et al. 2012). Neurons were transfected with either GFP- or Halo-tagged St3 and α4β2R subunit cDNAs, with β2 subunits tagged with an extracellular HA epitope to allow for assaying α4β2R surface expression (Fig. 2A). We found that a high percentage of neurons transfected with the St3 construct were also transfected with the α4β2R subunit cDNAs. As others have observed (Thayer, Jan et al. 2013), the Golgi in the somata of cultured pyramidal neurons is much larger and more extensive than the Golgi in other mammalian cells, such as in HEK cells (Fig. 2A, C, and E).

**Figure 2.**
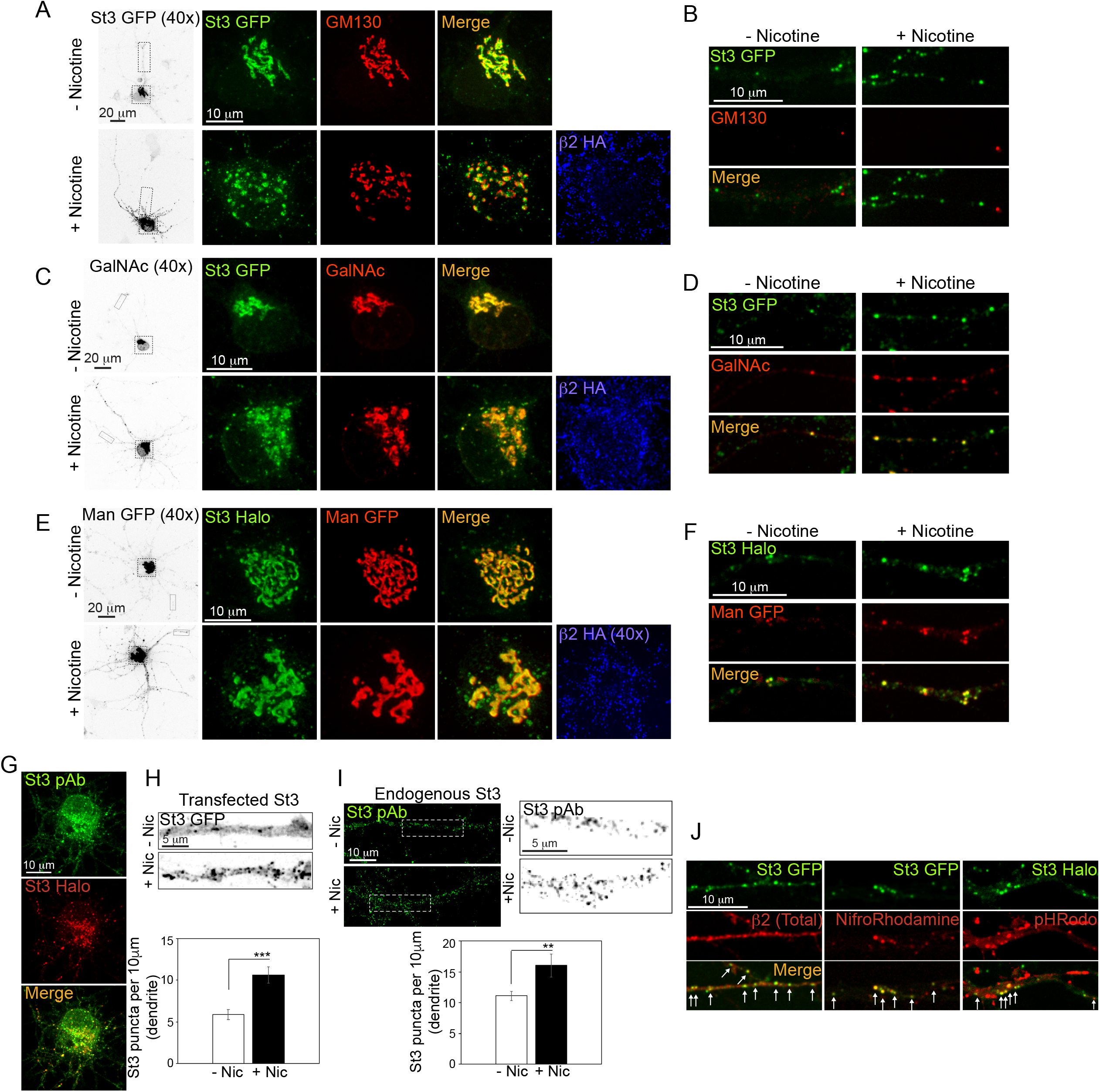
Nicotine-induced Golgi dispersal in primary cortical culture neurons. A. Distribution of St3 in cultured neurons. DIV 10 neurons (E18 rat pups) were transfected with St3-GFP (green) and nicotinic receptor subunits α4 and β2HA for 24 hours, and treated with or without 1μM nicotine for 17 hours. Cultures were fixed and immunostained for GM130 (red; secondary antibody anti-mouse Alexa 568). Inverted monochrome images of neurons expressing St3-GFP taken with 40x objective (left). Nicotine-induced distribution of larger GM130/St3 structures and smaller St3-only vesicles in the somata of neurons expressing α4β2HA (anti HA; blue) B. Dendritic processes showing distribution of St3-GFP. Nicotine increased the density of St3-GFP vesicles in dendrites. C. Co-distribution of St3-GFP and GalNAc T2-mCherry in cultured neurons. Cultures were transfected with St3-GFP (green), GalNAc T2-mCherry (red) and α4β2HA (anti-HA; blue) and treated with 1μM nicotine for 17 hours. Inverted monochrome images of neurons expressing GalNAc T2-mCherry (40x objective; left). St3-GFP and GalNAc T2 co-label intact Golgi structures. GalNAc T2 co-distributes to only a subset of St3-containing vesicles in the somata. D. Nicotine-induced increase in the distribution of GalNAc T2 in the dendrites of cultured neurons. St3-GFP overlaps with the majority of GalNAc T2-containing vesicles in the dendrites. E. Cultures were transfected with St3-Halo (green), α-Mannosidase II-GFP (Man-GFP; red) and α4β2HA (anti-HA; blue) and treated with or without 1μM nicotine for 17 hours. Inverted monochrome images of neurons expressing Man-GFP (40x objective; left). In the somata, Man-GFP co-distributed with St3-Halo in cis/medial Golgi structures, with very little localization to the small St3 vesicles. F. Nicotine increased the number of Golgi vesicles in dendrites that show overlap of Man-GFP and St3-Halo. G. Specificity of St3 polyclonal antibody (St3 pAb). Cultures were transfected with St3-Halo for 24 hours, then fixed and stained with anti-St3 antibody (St3 pAb). The St3 pAb labeled the majority of the St3-Halo vesicles in the somata and dendrites. H. Quantification of transfected St3 in dendrites. Cultures were transfected with St3-GFP and α4 and β2 nicotinic receptor subunits. Inverted monochromatic images of dendrites expressing St3-GFP (top). Scale bar, 5 μm. The number of St3-GFP puncta per 10 μm was quantified, data are shown as mean ± SEM, control cells, 5.9 ± 0.6; nicotine cells, 10.6 ± 1.0 (n = 9 neurons and 18 dendrites per group, ***p<0.000001). Detection of endogenous St3 in dendrites using polyclonal anti-St3 antibody (St3 pAb; left). Scale bar, 10 μm. Inset showing inverted monochromatic image (right). Scale bar, 5 μm. Quantification of the number of puncta recognized by St3 pAb per 10 μm, data are shown as mean ± SEM, control cells, 11.1 ± 0.8; nicotine cells, 16.1 ± 1.9 (n = 8-11 neurons per group, **p<0.008). J. High affinity α4β2 nicotinic receptors are present in acidic St3 vesicles. Cultures were transfected with either St3-GFP or St3-Halo (green) along with α4 and β2HA subunits. Left panel shows total β2HA distribution after permeabilization, fixation and staining with anti-HA (red). β2 subunits showed a reticulated distribution in the endoplasmic reticulum. It also co-localized with St3-GFP vesicles in the dendrites. Middle panel shows the presence of α4β2 receptors in St3-GFP vesicles (green) following labeling with a high affinity nicotinic receptor ligand, nifrorhodamine (red). Right panel shows the acidic nature of a subset of St3-Halo-containing vesicles (green) after loading the neurons with a pH sensitive dye, pHrodo (red). Arrows represent vesicles where co-labeling was observed.

Two other features distinguish the somatic Golgi of cultured neurons from the Golgi apparatus in most mammalian cells. First, the somatic Golgi in neurons is generally more dispersed compared to HEK cells, in terms of the number of smaller St3-containing puncta that lack GM130 (Fig. 2A, top panel). This dispersal is independent of the presence of α4β2Rs and nicotine, and has not been well characterized because it is much less pronounced with cis-Golgi markers, such as GM130, which have been widely used to identify Golgi membranes in cultured mammalian neurons. A second distinctive feature of the somatic Golgi morphology of neurons is that Golgi dispersal is highly variable. This neuron-to-neuron variability is illustrated in Supplemental Fig. S2-1 and was first noted by Ramon y Cajal in 1914, in Golgi-stained cat dorsal root ganglion cells (Ramon y Cajal 1914). As described previously (Thayer, Jan et al. 2013), changes in somatic Golgi morphology arise because Golgi dispersal in neurons is dependent on neuronal excitability and synaptic activity. Thus, variability in somatic Golgi morphology is likely caused by the neuron-to-neuron variability of synaptic and electrical activity in the cultures. The variability in neuronal Golgi morphology, when compared to other cells, limited our ability to quantify changes with nicotine exposure, such as that performed on the HEK cells (Fig. 1D).

An example of how nicotine exposure alters the morphology of the somatic Golgi of cultured pyramidal neurons is displayed in Figs. 2A - F. As observed in HEK cells (Fig. 1A) changes in Golgi morphology in response to nicotine exposure were only observed in neurons expressing α4β2Rs. To ensure that the neurons we assayed were expressing α4β2Rs, we transfected neurons with α4 and HA-tagged β2 subunits, which allowed us to perform live staining for cell-surface β2 subunits with anti-HA Abs (Figs. 2A, C, E, right panels). On the top of each panel are images from untreated neurons, and on the bottom, images for neurons treated with nicotine. On the left are images of the whole neuron with boxed areas for the soma (on the right) and examples of single dendrites (Fig. 2B, D and F) from each neuron. Neurons were transfected with α4β2R subunits, fluorescently tagged St3 or other Golgi markers: GM130 in Fig. 2A, B, GalNAcT2 in Fig. 2C, D or mannosidase II (Man-GFP) in Fig. 2E, F. The Golgi distribution of GalNAcT2 and Man is relatively more medial than that of the trans-Golgi St3 or cis-Golgi GM130 (Table 1). These distributions explain why St3 and GM130 staining patterns in the somatic Golgi (Fig. 2A) appear more polarized compared to St3 and GalNAcT2 (Fig. 2C) or Man (Fig. 2E). Similar to HEK cells, the somatic Golgi was well labeled by the GM130 Ab, and we observed two types of structures (Fig. 2A). The larger structures contained both GM130 and St3, with a polarized distribution typical of the GA, whereas the smaller structures contained only St3. There was some overlap of GalNAcT2 (Fig. 2C) and Man (Fig. 2E) with St3 staining in the smaller Golgi membranes, but many of these structures appeared to contain only St3.

The distribution of Golgi in the dendrites was different from that in the somata. Virtually all of the dendritic structures were similar in size to the smaller puncta in somata containing St3-GFP, and to a lesser degree, Man and GalNac. Many of these smaller puncta were mobile (Supplemental Fig. S2-4 video) and appear similar to “Golgi satellites” previously described (Mikhaylova, Bera et al. 2016). GalNAcT2 (Fig. 2D) and Man (Fig. 2F) were observed in most of the smaller St3-containing puncta, and nicotine increased the number of puncta labeled by all three Golgi markers. Only a small fraction of Golgi structures in dendrites were labeled by GM130 Ab, which correspond to the “Golgi outposts” (Hanus and Ehlers 2016) previously defined largely by GM130 staining. The number of GM130-stained Golgi membranes was not increased significantly by nicotine (Fig. 2A). Taken together, these findings are in contrast to observations using Golgi markers that are structural proteins, such as GM130, from which it has been concluded that there is very little functional Golgi in dendrites (Hanus and Ehlers 2016).

Because of the variability of the somatic Golgi morphology from neuron to neuron (Supplemental Fig. S2) and the difficulty in identifying the same neuron before and after nicotine exposure, nicotine-induced changes in somata were not always readily apparent (Fig. 2E). However, in the dendrites, nicotine-induced changes were more uniform. To quantify changes in dendrites, we examined the number and density of St3-containing puncta using both overexpression of the St3-GFP construct and a St3-specific Ab to assay endogenous protein (Fig. 2H and I). The St3-specific Ab staining overlapped significantly with St3-Halo expression, thus validating the specificity of the St3 Ab (Fig. 2G). Estimates of St3 punctal density (puncta/10 μm) were determined on parallel cultures (Fig. 2H, I; Table II). The punctal density estimate using the St3-specific Ab is ∼2-fold higher than the estimate using St3-GFP expression, consistent with St3-GFP only being incorporated in about 50% of the available St3-containing structures. Nicotine exposure resulted in significant increases in punctal density using both approaches. The increase estimated by analyzing St3-GFP expression was slightly higher than that of endogenous St3 (1.8-fold, versus 1.4-fold). This difference may be due to the fact that St3-GFP expression and trafficking largely occurred during the period of nicotine exposure (∼17 hours), which may not be true for endogenous St3 expression and trafficking.

**Table 2.**
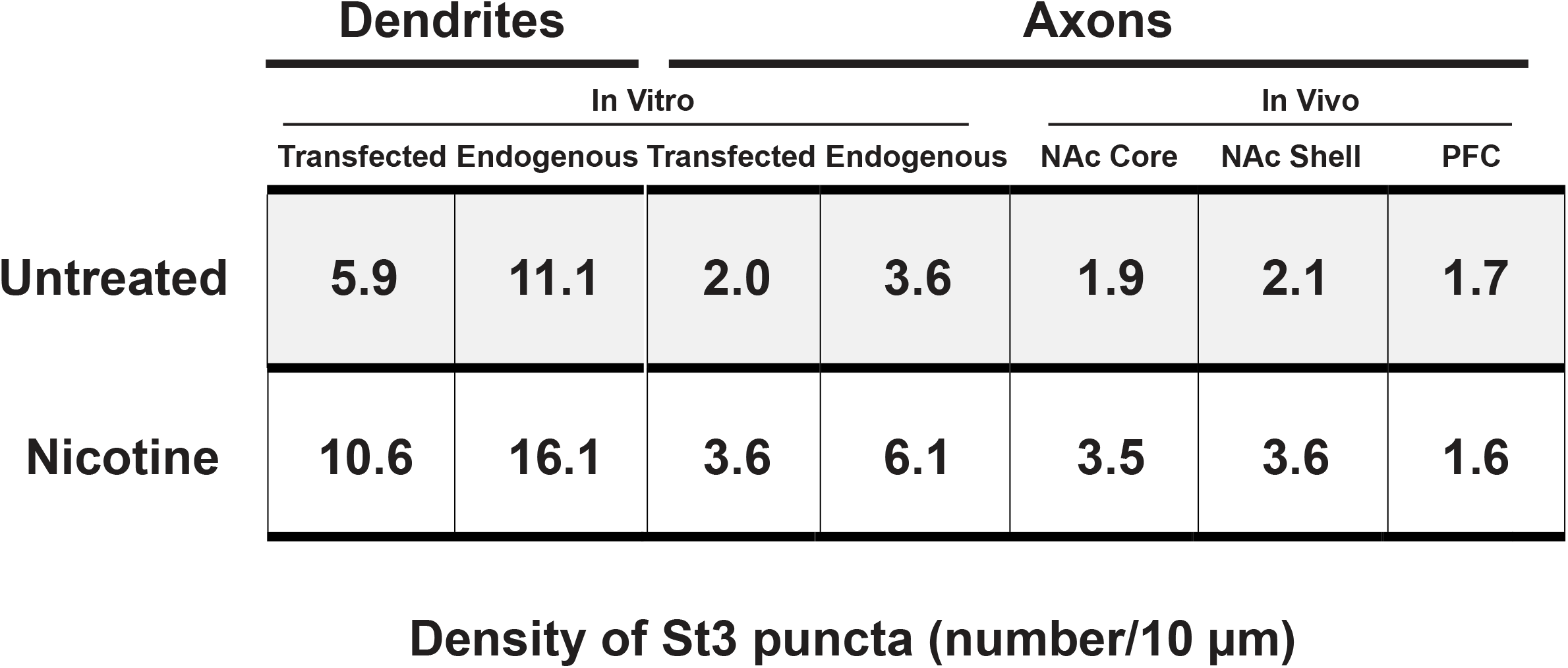
Comparison of *in vitro* and *in vivo* St3 punctal densities in neuronal processes.

We then assessed whether St3-containing dispersed Golgi membranes in neurons shared features of the α4β2R-containing acidic vesicles (Govind, Vallejo et al. 2017). Similar to HEK cells (Fig. 1G), nifrorhodamine stained a subset of the small St3-labeled puncta in dendrites (Fig. 2J, middle panels) and somata (Supplemental Fig. S2-2A), but not the larger Golgi membranes marked by both St3 and GM130 (Fig. 2A). Shown in supplemental Fig. S2-2 (top and bottom panels) is a comparison of nifrorhodamine staining with that of HA-tagged β2 subunits in neurons. The staining for β2 subunits in neurons overexpressing α4β2Rs appears to have little overlap nifrorhodamine labeling (sFig. 2A, B). The reason for this lack of overlap in these cultured neurons is that only a fraction of total β2 subunits assemble into α4β2Rs (Govind, Walsh et al. 2012) and even a smaller fraction of the assemble receptors contain high-affinity nicotine binding sites that bind nifrorhodamine. β2 subunit staining has a reticulated pattern in the soma and stains the nuclear membrane, consistent with an ER distribution, which is where unassembled β2 subunits assemble into α4β2Rs. Surprisingly, there is little or no nifrorhodamine staining of β2 subunits in the ER or the large St3-containing Golgi structures. With respect to intracellular organelles, we only observe nifrorhodamine labeling in the small ST3 puncta. This finding suggests that high-affinity nicotine binding sites form on α4β2Rs when they are trafficked to the small, St3-containing Golgi puncta, or just before entry into that compartment. In dendrites (Fig. 2J), the ER is relatively less extensive when compared to somata and, similarly, the amount of β2 subunit staining in the ER less extensive. This allowed us to compare the distribution of β2 subunits (Fig. 2J, left panel), nifrorhodamine (middle panel) and pHrodo (right panel) with St3. All three partially overlapped with the small St3-containing puncta, establishing that high-affinity α4β2Rs are found in a subset of the St3-containing dispersed Golgi membranes.

### Dispersed Golgi membranes in axons and effects of nicotine

Golgi structures have not been observed in axons of mature neurons using standard Golgi markers such as GM130 (Hanus and Ehlers 2016). However, early in axon formation they are present and reappear during axon regeneration (Merianda, Lin et al. 2009). Additional studies suggest that Golgi membranes are present in the axons of mature neurons. In cultures, the trans-Golgi network (TGN) protein, TGN38, is present in axonal mobile vesicles (Nakata, Terada et al. 1998) and the medial- to trans-Golgi protein, ST3 Beta-Galactoside alpha-2,3-Sialyltransferase 5 (St3Gal5), is also observed in axons (Stern and Tiemeyer 2001). St3Gal5 adds sialic acid to glycolipids and is distinct from the St3 Golgi enzyme assayed in Figs. 1 and 2, which adds sialic acid to protein N-linked glycans. More recently, standard Golgi markers were found at nodes of Ranvier of myelinated axons (Gonzalez, Canovas et al. 2016). Since the Golgi markers used in Fig. 2 were much more abundant in dendrites than standard Golgi markers, we tested for their presence in axons.

Axons were distinguished from dendrites by their more uniform, narrow diameter, and by antibody staining for neurofilament heavy chain (NFH; Fig. 3A, right). We observed no GM130 in axons (Fig. 3A), confirming previous findings in cultured neurons from which it was concluded that axons lack any functional Golgi membranes (Hanus and Ehlers 2016). In contrast to GM130 Ab staining, labeling of endogenous St3 (Fig. 3B) and transfection of tagged versions of St3, overlapped with NFH Ab staining (Fig. 3C). We tested for the axonal presence of two other Golgi enzymes, Mannosidase II and St3Gal5, by expressing Man-GFP or labeling for endogenous St3Gal5 using an Ab that previously stained axons of cultured neurons (Stern and Tiemeyer 2001). Both Golgi resident proteins exhibited overlap with St3-(Supplemental Fig. S3-1A and B) and β2-containing puncta (Supplemental Fig. S3-1C) in axons, and many St3-containing puncta were mobile (Supplemental Fig. S3-2; video), moving at a rate of ∼1 μm/sec. Nicotine exposure caused an increase in the number of St3 puncta (Fig. 3B and C). Punctal densities of endogenous and transfected St3 (St3-Halo) were quantified and compared with those obtained in dendrites (Table II). As we observed in dendrites, endogenous St3 punctal density was almost 2-fold greater when compared to transfected St3, visualized by HaloTag ligand labeling (Table III). These values were ∼3-fold higher in dendrites versus axons for both endogenous and transfected St3. Using a Venus cell-fill, we measured the cross-section of dendrites in our cultures and found it is ∼3-fold greater than for axons, suggesting that St3 punctal density is similar in both compartments when normalized to their respective diameters.

**Figure 3.**
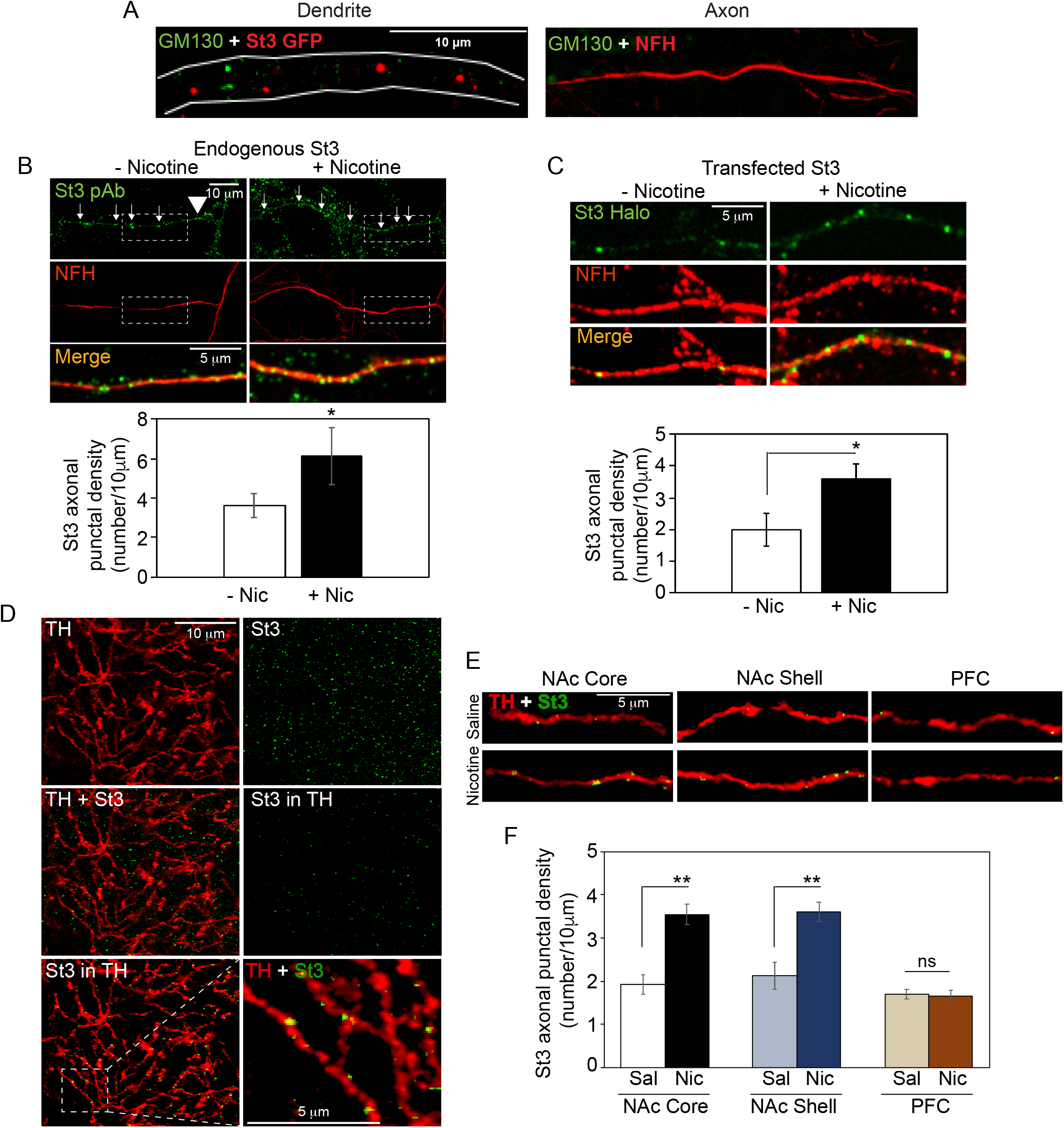
Nicotine-induced increase in axonal Golgi membrane density *in vitro* and *in vivo*. A. DIV 10 neurons were transfected with St3-GFP (red) for 24 hours, fixed and immunostained with a mAb against the conventional *cis*-Golgi marker, GM130 (green). GM130 labeling exhibited a punctate dendritic distribution (left panel) and a lack of axonal labeling (right panel). Scale bar, 10 μm. B. Detection of endogenous St3 in axons using St3 pAb (top panels). Axons were identified by staining with an Ab against the axonal marker, anti-NFH (middle panels). Cultures were transfected with nicotinic receptor subunits α4 and β2HA for 24 hours and treated with or without 1μM nicotine for 17 hours. The number of endogenous St3 puncta per 10 μm was quantified, and nicotine treatment resulted in a significant increase in St3 density in axons. Data are shown as mean ± SEM, control cells, 3.6 ± 0.6; nicotine cells, 6.1 ± 1.4 (n = 5 fields per group, *p<0.03 relative to control). Scale bar, 10 μm. C. Detection of transfected St3 in axons. Cultures were transfected with St3-Halo (green) and α4β2HA, treated with or without 1μM nicotine for 17 hours, and fixed and immunostained with anti-NFH (red). Axonal density of transfected St3 was increased in nicotine-treated cultures. Data are shown as mean ± SEM, control cells, 1.9 ± 0.5; nicotine cells, 3.6 ± 0.4 (n = 5 fields per group, *p<0.02 relative to control). Scale bar, 5 μm. D. Golgi marker enzymes are localized to dopaminergic terminals in nucleus accumbens. Top panels, double immunolabeling of St3 (green) and dopaminergic fibers (anti-TH; red) in the nucleus accumbens (NAc) from mouse brain. Scale bar, 10 μm. Middle panels, overlay of TH and St3 channels (left) and isolation of St3 puncta localized within the dopaminergic fibers (right). Bottom panels, low (left) and high (right; inset of left panel) magnification examples of St3 puncta localized within dopaminergic terminals of the NAc. Inset scale bar, 5 μm. E. Representative micrographs displaying isolated St3 puncta within dopaminergic terminals of the NAc core and shell regions, and the prefrontal cortex (PFC). Mice received either 100 µg/ml nicotine via drinking water, or regular water, daily for 3 weeks. Scale bar, 5 μm. F. Quantification of St3 punctal density within dopaminergic terminals displayed in E. Nicotine exposure significantly increased levels of St3 in NAc core and shell, but had no effect on St3 density in dopaminergic terminals of the PFC. Data are shown as mean ± SEM, NAc core, control, 1.9 ± 0.2, nicotine, 3.5 ± 0.2; NAc shell, control, 2.1 ± 0.3, nicotine, 3.6 ± 0.2; PFC, control, 1.7 ± 0.1, nicotine, 1.6 ± 0.1; obtained from 3 vehicle-treated and 4 nicotine-treated mice; measurements obtained in 3 sub-regions in 3 separate volumes per region/per mouse (core, **p<0.002, compared to vehicle; shell, **p<0.003, compared to vehicle).

**Table 3.**
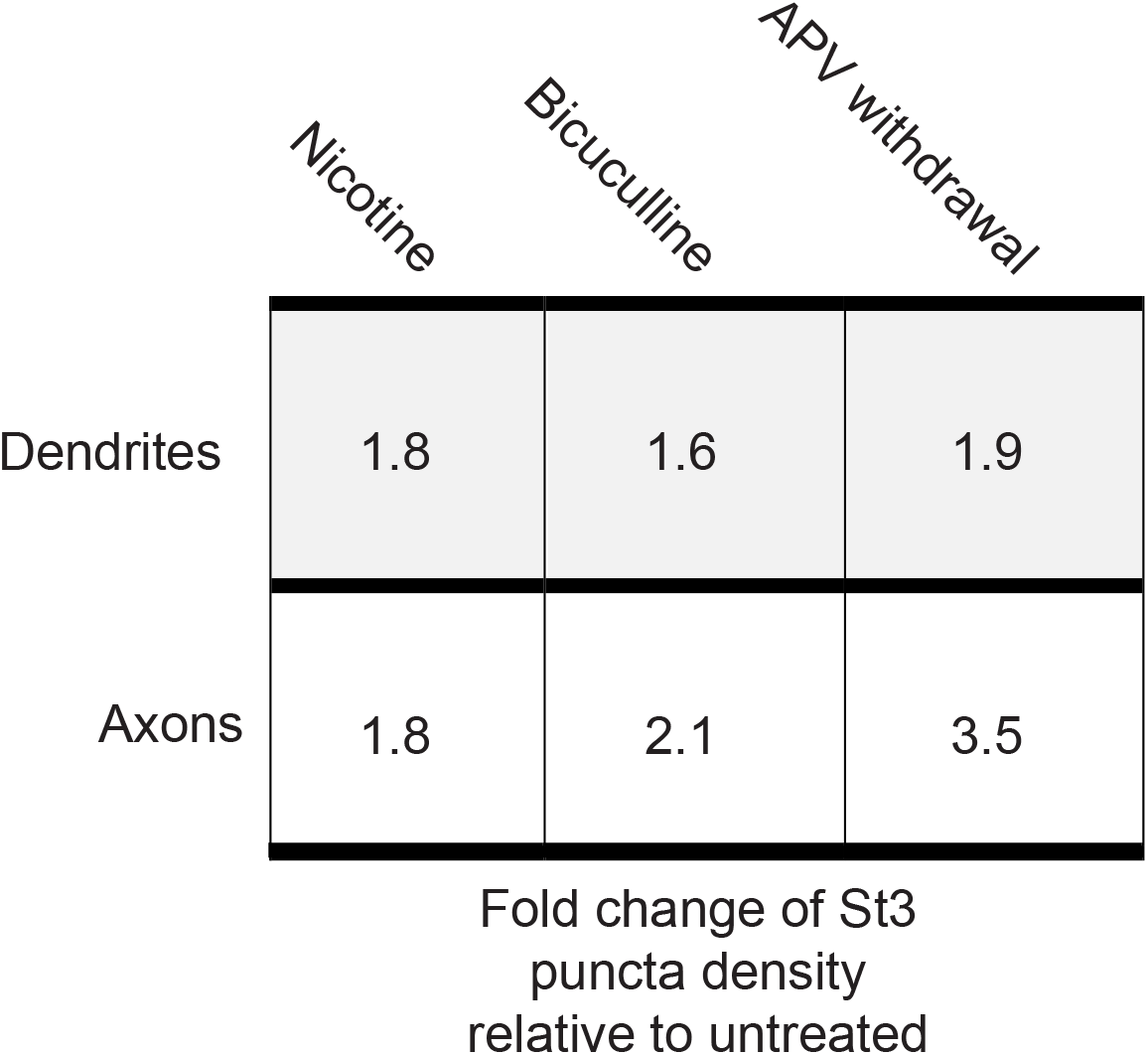
Activity-dependent changes in St3 punctal density in dendrites and axons of primary cortical cultures.

Previously, Tarren et al., 2017, used a YFP-tagged α4 subunit knock-in mouse to demonstrate that α4 subunit-containing puncta were localized to terminals of ventral tegmental area (VTA) dopaminergic neurons identified by tyrosine hydroxylase (TH) staining (Tarren, Lester et al. 2017). Mice that were administered ethanol exhibited increased α4 subunit punctal densities at VTA dopaminergic neuron terminals in the nucleus accumbens (NAc) and amygdala, but not at other terminals such as in the prefrontal cortex (PFC). We performed similar experiments to test whether St3-containing puncta are found in these terminals, in a pattern similar to the α4 subunit puncta, and also tested whether nicotine exposure increased their number. We identified VTA dopaminergic neuron terminals with TH-specific Abs and co-stained with the St3 Ab. We observed many St3-containing puncta throughout each region we analyzed. We also found that a subset of the puncta overlapped with TH Ab staining. An example of the total St3 staining (green puncta) is displayed in Fig. 3D (top, right panel), as well as the masking and isolation of St3 labeling within TH-positive membranes, in order to examine St3 puncta found in the dopaminergic terminals. This analysis revealed a significant number of St3 puncta within dopaminergic neuron terminals (Fig. 3D middle, right panel; Bottom panels), similar to the α4 subunit puncta previously observed using YFP-α4 knock-in mice. The dopaminergic axonal punctal densities measured in three different regions, NAc core, NAc shell and PFC were similar, but approximately half of the density measured for the cultures in vitro (Table II).

Administration of nicotine in the drinking water of mice for 3 weeks resulted in increased axonal St3 punctal number in NAc core and shell, but not in the PFC, when compared to untreated mice (Fig. 3E and F). These results correspond to the increased numbers of YFP-α4 puncta observed following alcohol exposure in YFP-α4 knock-in mice (Tarren, Lester et al. 2017). There, increases also occurred in VTA dopaminergic neuron terminals in the NAc core and shell, but not in the PFC. The increase in St3 puncta we measured in NAc core and shell were comparable to what we observed in vitro, in the axons and dendrites of cultured neurons (Table II). This provides strong evidence that nicotine-induced Golgi dispersal occurs in vivo, in VTA dopaminergic neurons, and suggests that dispersed Golgi elements may contribute to nicotine’s addictive effects there.

### Changes in synaptic activity regulate GA dispersal and distribution

In this study, we observed that long-term nicotine exposure *in vitro* and *in vivo* induces a dispersal and redistribution of the Golgi apparatus, but only in neurons expressing α4β2Rs. We have also found that the degree to which the somatic Golgi of cultured neurons is dispersed is highly variable (Supplemental Fig. S2-1). Somatic Golgi dispersal in neurons has been shown to be induced by increasing levels of excitability and synaptic activity (Thayer, Jan et al. 2013), but activity-induced Golgi dispersal in neurites has not been analyzed using other Golgi markers. Therefore, we tested whether drug treatments known to disperse the somatic Golgi also altered levels of Golgi membranes in dendrites and axons. Neurons were transfected with St3-Halo and Venus, to fill the cytoplasm and allow visualization of processes (Fig. 4A, C). Cortical cultures were stimulated by applying the GABA_A_ receptor antagonist bicuculline for 18-20 h, or by withdrawing the NMDA receptor antagonist APV for 18-20 h following 2 days of APV treatment. Bicuculline treatment led to a 1.5-fold increase in St3-Halo-positive Golgi vesicles in dendrites and a 2-fold increase in the axons of pyramidal neurons, compared to vehicle treatment (Fig. 4B), and NMDA receptor activation via APV withdrawal led to a similar effect compared to vehicle or chronic APV treatment (Fig. 4D). The St3-Halo punctal density in these experiments was lower than what was observed in neurons transfected with α4β2Rs, but this is likely an artifact of co-transfection with Venus. Our results demonstrate that increasing synaptic activity promotes Golgi dispersal in neurons and increases the trafficking and levels of Golgi membranes in dendrites and axons in a manner similar to nicotine exposure, but independent of α4β2R expression. These activity-regulated dendritic and axonal Golgi compartments may function in protein secretion at local subdomains and contribute to the molecular mechanisms underlying synaptic plasticity.

**Figure 4.**
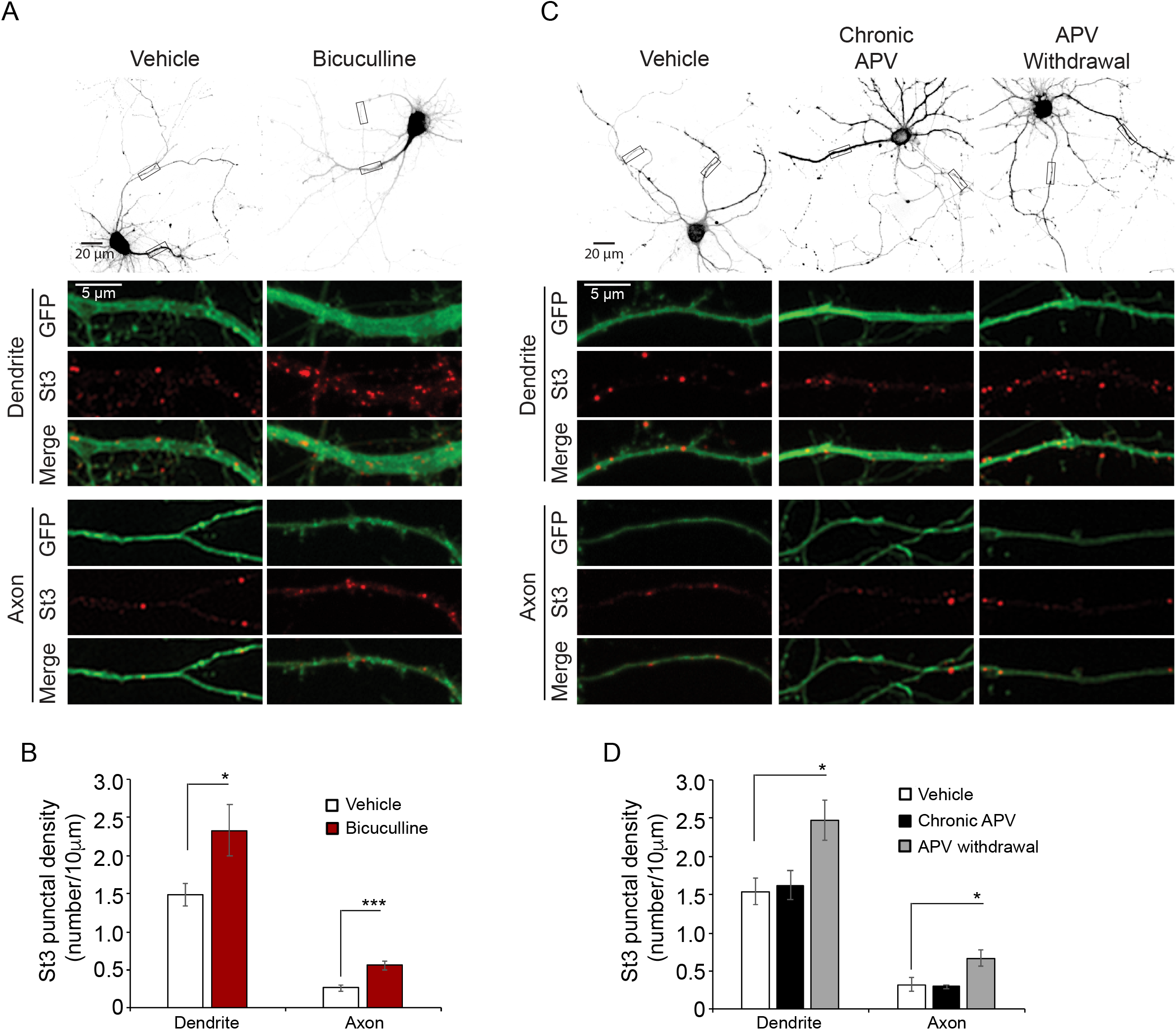
A. Activity-induced increase in Golgi membrane density. Top, 40× magnification grayscale images of representative GFP-expressing neurons subjected to vehicle (left) or bicuculline (right) treatment and used for quantification of St3-Halo-positive puncta. Rectangles indicate the location of the 100× magnification insets showing dendrites (middle three rows) and axons (lower three rows). B. Dendrites (left) and axons (right) from neurons treated with bicuculline displayed significantly higher densities of St3-positive puncta than those treated with either vehicle. Data are shown as mean ± SEM, Bicuculline, dendrites: control cells, 1.5 ± 0.2; bicuculline cells, 2.3 ± 0.3 (n = 12 neurons per group, *p<0.05); axons: control cells, 0.27 ± 0.04; bicuculline cells, 0.56 ± 0.06 (n = 12 neurons per group, ***p<0.0005). C. Top, 40× magnification grayscale images of representative GFP-expressing neurons subjected to vehicle (left), chronic APV (center), or APV withdrawal (right) treatment and used for quantification of St3-Halo-positive puncta. Rectangles indicate the location of the 100× magnification insets showing dendrites (middle three rows) and axons (lower three rows). D. Dendrites (left) and axons (right) from neurons subjected to APV withdrawal displayed significantly higher densities of St3 positive puncta than those treated with either vehicle or chronic APV. Data are shown as mean ± SEM, APV withdrawal, dendrites: control cells, 1.5 ± 0.2; chronic APV cells, 1.6 ± 0.2; APV withdrawal cells, 2.5 ± 0.3 (n = 7 neurons per group, *p<0.02); axons: control cells, 0.32 ± 0.1; chronic APV cells, 0.30 ± 0.03; APV withdrawal cells, 0.67 ± 0.12 (n = 7 neurons per group, *p<0.02).

### GA dispersal by nicotine is similar to GA dispersal by nocodazole

To test how nicotine and changes in synaptic activity result in dispersal of the Golgi, we examined the processes involved in dispersal in more detail. There appear to be different mechanisms by which Golgi dispersal occurs. In certain types of neurodegenerative disorders, such as ALS, Parkinson’s Disease, and Alzheimer’s Disease, Golgi dispersal or “fragmentation” is thought to be similar to the dispersal that occurs during apoptosis, where the Golgi undergoes fission and stack uncoupling events that cause it to break apart or fragment (Mancini, Machamer et al. 2000). This type of dispersal results in deficits in GA function and is irreversible. In contrast, if microtubule polymerization is blocked by nocodazole, Golgi fragmentation occurs by a different process. After nocodazole treatment, the balance between forward trafficking through the Golgi from the ER and retro-directed trafficking back to the ER is altered due to a loss of microtubule-dependent forward trafficking. This imbalance causes the preexisting GA to diminish and the forward trafficking to reroute through smaller Golgi “ministacks’ that arise at more peripheral ER exit sites (ERES; (Cole, Sciaky et al. 1996, Storrie, White et al. 1998). Nocodazole-induced ministacks are polarized, just like the larger ministack membranes containing both St3 and GM130 we observed with nicotine-induced dispersal (Figs. 1C, D and E).

To examine how GA dispersal by nicotine occurs, we compared GA dispersal induced by nocodazole and nicotine in α4β2R-expressing HEK cells and primary cortical cultures (Fig. 5). By eye, we could not distinguish the dispersed Golgi elements that resulted from either drug alone, or co-administered together in the HEK cells (Fig. 5A). Similar to nicotine treatment, nocadozole resulted in two types of dispersed Golgi membranes: larger polarized “ministacks” containing St3-GFP and GM130, and smaller St3-only fragments, both of which had the same size distribution (Fig. 5B and Supplemental Fig. S5-1A). Using the same quantitative approach we employed to analyze nicotine-induced dispersal (Fig. 1D), we found only small differences in how each treatment increased the numbers of the two types of membranes. Nocadozole treatment led to greater numbers of the polarized ministacks (Fig 5B; top;**p= 0.0002 for Noc; *p= 0.03 for Nic), whereas nicotine treatment resulted in slightly more of the smaller, St3-only fragments (Fig. 5B; bottom; **p= 0.00004 for Nic; *p= 0.003 for Noc). The most significant difference between nicotine and nocadozole treatments were in their effects on microtubule distribution in HEK cells (Fig. 5C). As expected, nocadozole treatment highly reduced the microtubule content in HEK cells, as assayed by co-expression of an mCherry-tagged, microtubule-associated protein, Ensconsin (Fig. 5C). Nicotine had no apparent effects on microtubule distribution, indicating that nicotine-induced GA dispersal occurs through a mechanism different from nocadozole.

**Figure 5.**
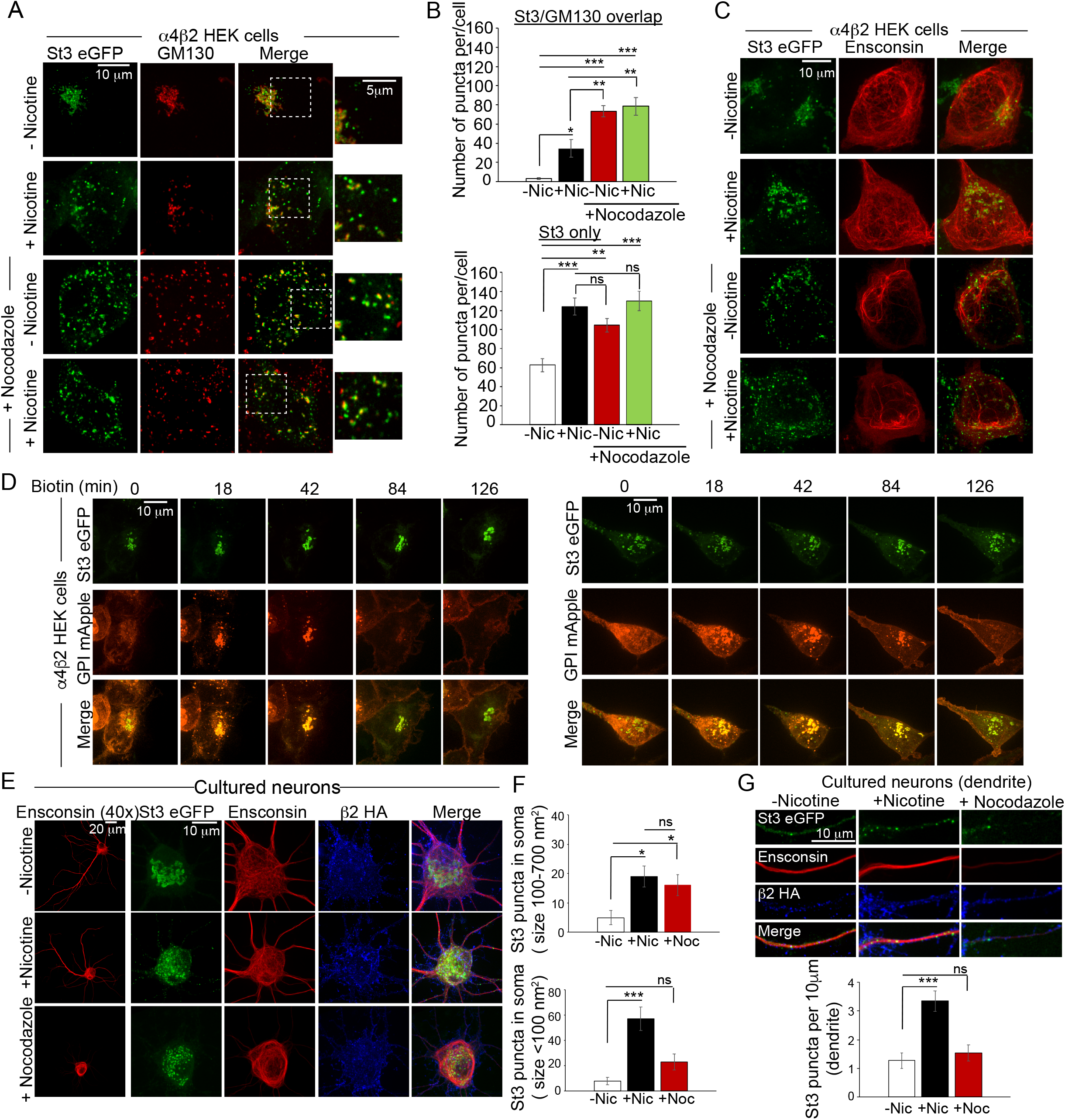
Dispersed Golgi elements are functional. A. Similarities and differences between Golgi dispersal induced by nicotine and nocodazole. α4β2 HEK cells were transfected with St3-GFP (green). Cells were treated with 10 μM nicotine for 17 hours and further treated for 4 hours with 25 μM nocodazole. Cells were fixed, permeabilized and immunostained with anti GM130 antibody (red). Scale bar, 10 μm. Inset scale bar, 5 μm. B. Quantification of the number of puncta per cell displaying St3/GM130 overlap (top) or St3 only (bottom). Data are shown as mean ± SEM, for St3/GM130 overlap, control cells, 3.1 ± 1.0; nicotine cells, 34.6 ± 9.0; nocodazole cells, 73.3 ± 6.0; nicotine and nocodazole cells, 78.5 ± 9.3 (n = 10 cells per group, con vs nic, *p<0.035; con vs noc, ***p<0.00003; con vs nic + noc, ***p<0.000003; nic vs noc, p<0.007; nic vs nic + noc, p<0.003) and for St3 only, control cells, 62.6 ± 6.9; nicotine cells, 124.1 ± 8.9; nocodazole cells, 104.5 ± 6.9; nicotine and nocodazole cells, 130 ± 10 (n = 10 cells per group, con vs nic, ***p<0.00004; con vs noc, **p<0.004; con vs nic + noc, ***p<0.000005). C. Effect of nicotine on microtubule stability. α4β2 HEK cells were transfected with St3-GFP (green) and the microtubule binding protein, Ensconsin-mCherry (red). Cells were treated with nicotine and nocodazole as in A. D. α4β2 HEK cells were transfected with St3-GFP and RUSH-GPI-mApple (retained in the endoplasmic reticulum via an ER targeted streptavidin/streptavidin binding peptide hook) for 24 hours and treated with 10 μM nicotine for 17 hours. Left panels, live imaging of cells showing transport of GPI-mApple to plasma membrane occuring via intact Golgi following biotin-mediated release from ER. Right panels, GPI-mApple traffics through dispersed Golgi elements following release from ER (right). E. Comparison of the effect of nicotine and nocodazole on primary cortical cultures. Cultures of neurons (DIV 10) were transfected with α4, β2HA, St3-GFP and Ensconsin-mCherry for 24 hours. Neurons were treated with 1 μM nicotine for 17 hours and further treated with 8 μM nocodazole for 4 hours. Neuronal cultures were live labeled with anti-HA antibody (β2HA; blue), fixed and stained with secondary anti-mouse Alexa 647. Left panels, 40x images of neurons showing Ensconsin-mCherry (red) distribution in somata and dendrites. Scale bar, 20 μm. Right panels, St3-GFP (green) appeared dispersed in the somata of nicotine and nocodazole treated cultures, compared to untreated cultures. Scale bar, 10 μm. F. Quantification of the number of larger (100-700 nm^2^; top) and smaller (less than 100 nm^2^; bottom) St3-GFP puncta. Top, data are shown as mean ± SEM, control cells, 5.3 ± 2.4; nicotine cells, 18.8 ± 3.5; nocodazole cells, 16.4 ± 3.6 (n = 6-8 cells per group, con vs nic, *p<0.02; con vs noc, *p<0.03; nic vs noc, ns). Bottom, data are shown as mean ± SEM, control cells, 8.2 ± 2.9; nicotine cells, 56.6 ± 9.3; nocodazole cells, 22.7 ± 6.2 (n = 6-8 cells per group, con vs nic, ***p<0.0004; con vs noc, ns). G. Dendritic distribution of St3-GFP in neurons from E after treatment with nicotine and nocodazole. Scale bar, 10 μm. G. Quantification of the number of St3-GFP puncta per 10 μm of dendrite (bottom). Data are shown as mean ± SEM, control cells, 1.3 ± 0.3; nicotine cells, 3.3 ± 0.3; nocodazole cells, 1.5 ± 0.3 (n = 6-8 cells per group, con vs nic, ***p<0.0006; con vs noc, ns).

As seen with nocodazole-induced dispersal of the GA (Fourriere, Divoux et al. 2016), we find that the dispersed membranes induced by nicotine are functional (Fig. 5D) and that the dispersal itself is reversible (Supplemental Figs. S5-2B bottom panel; S5-2C). We used the retention using selective hooks (RUSH) system (Boncompain, Divoux et al. 2012) to test whether cargo trafficked through the dispersed Golgi elements after nicotine treatment (Fig. 5D). The RUSH system can be used to hold fluorescent cargo in the ER until biotin is added to the cells, which releases the cargo from the ER by breaking a bond between an ER resident protein and the cargo. Here, the cargo was glycosylphosphatidylinositol fused to fluorescent mApple (GPI-mApple) and the ER-resident protein was KDEL-streptavidin. At the time biotin was added (t = 0), the St3-GFP distribution in α4β2R-expressing HEK cells indicates an intact GA while the GPI-mApple cargo is initially held in the ER (Fig 5D, left panels). Within 18 minutes following the addition of biotin, much of the GPI-mApple moved from the ER into the GA. This trafficking from the ER continued for 30 - 60 minutes, followed by trafficking from the GA to the plasma membrane. In the right panels is a single α4β2R-expressing HEK cell transfected with GPI-mApple cargo and St3-GFP, but exhibiting a dispersed GA following nicotine treatment. Despite the dispersed GA, GPI-mApple cargo still traffics through the dispersed membranes and then to the plasma membrane with roughly the same time-course as with the intact GA on the left.

GA dispersal in α4β2R-expressing HEK cells slowly reverses ∼22 to 48 hours after nicotine is removed from the medium (Supplemental Fig. S5-2B and C). This rate is slower than that of nicotine-induced dispersal, which occurs ∼2 to 4 hours after nicotine exposure (Supplemental Figs. S5-2A and S5-2B; top), and is somewhat slower than the rate of dispersal resulting from nocadozole treatment (Fourriere, Divoux et al. 2016). The rates at which dispersal by nicotine occur were similar to the rate of nicotine upregulation of α4β2Rs for the fast phase of the upregulation (Govind, Walsh et al. 2012), suggesting that Golgi dispersal with nicotine exposure occurs in parallel with receptor upregulation. Therefore, we used a radioactive ligand binding assay to test whether the nocadozole-induced Golgi dispersal itself could cause nicotine upregulation of α4β2Rs (Supplemental Fig. S5-1B). Nocadozole treatment of HEK cells did not result in an increase in the number of high-affinity α4β2R binding sites. However, treatment with nicotine and nocadozole together did result in a significant increase in binding site number. These findings suggest that Golgi dispersal in and of itself does not cause nicotine α4β2R upregulation, but dispersal induced by nocadozole significantly augments nicotine upregulation, suggesting that this process has a role in upregulation.

In the somata of cultured neurons, nocodazole- and nicotine-induced Golgi dispersal were similar, however microtubules were not affected by nicotine exposure (Fig. 5E and F). In dendrites (Fig. 5G), nocodazole did not cause the ∼2-fold increase in St3 punctal density we observed with nicotine treatment. One explanation for this might be that nocodazole-induced destabilization of microtubules in dendrites leads to a loss of vesicular trafficking from soma to dendrites along microtubules (Fig. 5G). However, we observed no significant increase in the number of small puncta (size <100nm^2^) in the somata, as would be expected if their transport were prevented by nocodazole treatment (Fig. 5F, bottom panel).

Another feature of nocodazole-induced Golgi dispersal is the close association between ERES and many of the dispersed Golgi elements (Cole, Sciaky et al. 1996, Storrie, White et al. 1998). Nocodazole causes dispersed Golgi to emerge from ERESs that are more peripheral relative to original ERES sites near the GA. A similar close association appears to occur with nicotine treatment in α4β2R-expressing HEK cells (Supplemental Fig. S5-3A) and in the somata of cultured neurons (Supplemental Fig. S5-3B), as shown by co-imaging St3 with either the ER (DsRed-ER) or ERESs (mCherry-Sec23). In dendrites, ERESs were found throughout the shaft (Supplemental Fig. S5-3C) and we observed a close association between them and St3-containing puncta (Supplemental Fig. S5-3D). In axons, ERESs were only observed in the axon initial segment (Supplemental Fig. S5-3D), and thus, the spatial alignment between the two largely does not occur. Close association between ERESs and dispersed Golgi membranes in dendrites is consistent with the possibility that a portion of the St3-containing structures emerge from ERESs there instead of being trafficked from the soma.

### Do secreted proteins traffic through dispersed Golgi membranes in dendrites and axons?

To gain more insight into how dispersed Golgi structures arrive in dendrites and axons, we tested whether fluorescent cargo held in the ER traffics to Golgi in the soma and then is transported to axons or dendrites. Alternatively, cargo could move directly from the local ER into the Golgi in dendrites and axons after ER release. To hold cargo in the ER, we first tried the same RUSH system as in Fig. 5D to examine GPI cargo trafficking through dendritic St3-containing puncta. Supplemental Fig. S6 displays an example of this cargo trafficking through the Golgi structures following release with biotin. However, there were technical difficulties applying this system to dendrites. First, there often appeared to be some “leak” between the ER and Golgi prior to release of the ER-bound cargo. In addition, there was a mobile pool of both the ER-bound cargo, which had a punctate distribution, and the St3-containing puncta, which made it difficult to establish when the cargo was trafficking through the Golgi puncta.

To counter these problems, we used an alternative system that allowed synchronized release of bound fluorescent cargo from the ER (Casler, Papanikou et al. 2019). In this system, the cargo, a modified DsRed-Express2, forms homo-aggregates in the ER lumen when expressed. The aggregates in the soma, dendrites and axons are displayed for a transfected cultured neuron before release (Fig. 6A). Upon addition of a membrane-permeable ligand, the cargo rapidly filled the ER lumen, as aggregates dissociate into tetramers (Fig. 6B; boxed dendrite in A). The cargo largely moved from the soma into the dendrite and reached a steady state as soon as we began imaging following ligand addition (4 minutes). Subsequently, the cargo in the ER lumen moved into static Man II-containing Golgi puncta along the dendrites. These data demonstrate the trafficking of ER cargo into the dispersed Golgi membranes in dendrites and are consistent with Golgi structures emerging from ERESs nearby.

**Figure 6.**
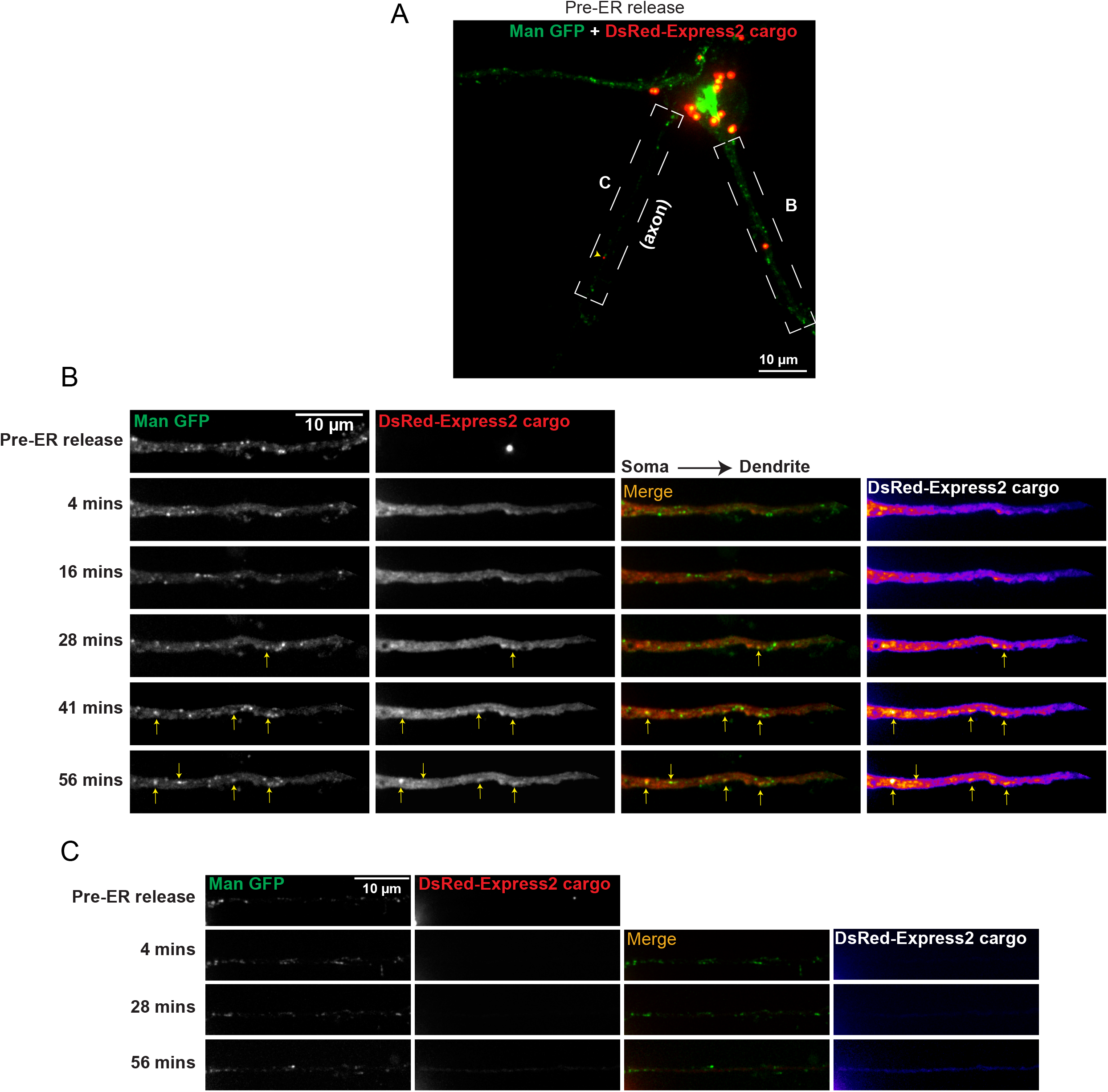
Secretory trafficking of cargo through dispersed Golgi membranes. A. DIV 10 primary cortical cultures were transfected for 24 hours with a bicistronic cDNA plasmid encoding Man-GFP (green) and a modified DsRed-Express2 (red) that forms aggregates in the ER lumen. Displayed is the whole neuron distribution of both secretory markers prior to ER release. Scale bar, 10 μm. B. Live-imaging of the boxed dendrite in A, pre-release and following addition of a synthetic ligand that dissolves the ER aggregates, allowing ER exit. Fluorescent cargo rapidly filled the ER lumen, then entered and accumulated in dispersed, static Golgi structures (Man-GFP, arrows). Image frames were acquired every 4 mins for 1 h, scale bar, 10 µm. C. Live-imaging of the boxed axon in A, pre-release and following addition of synthetic ligand. Man-GFP puncta were localized along the length of the axon, but axonal ER cargo trafficking was significantly delayed as compared to dendrites, taking almost 1 h to visualize. Scale bar, 10 µm.

The results in axons were very different from what we observed with trafficking in the dendrites (Fig. 6C). Unlike the dendrites, where the ER lumen was rapidly filled by cargo flow from the soma, cargo entry from soma to axon was extremely slow. There was little entry of the cargo even after ∼ 1 hour of imaging, suggesting that there is a diffusional barrier that slows the transport of ER lumenal cargo from the somatic to axonal ER. Such a structural feature is consistent with previous studies where diffusional barriers were found in the plasma membrane at the initial axon segment (Winckler, Forscher et al. 1999, Nakada, Ritchie et al. 2003). The extremely slow rate of cargo transport from the somatic ER to axonal ER prevented us from testing whether ER cargo exited locally from the ER and entered the Golgi membranes we observed in axons (Fig. 3).

## Discussion

### GA dispersal by nicotine and upregulation of α4β2Rs

Chronic nicotine exposure causes a set of long-term changes in α4β2Rs, which have been lumped together in a process called α4β2R upregulation. Nicotine upregulation of α4β2Rs was first measured as increases in α4β2R high-affinity binding site numbers in the brain, and later was found to correlate with increased α4β2R functional response after nicotine exposure (Marks, Burch et al. 1983, Schwartz and Kellar 1983, Benwell, Balfour et al. 1988, Breese, Adams et al. 1997). What causes α4β2R upregulation by nicotine and how it is involved in nicotine addiction are questions that remain largely unanswered. Our discovery of the dispersal of the GA by nicotine exposure and the involvement of α4β2Rs in the process provides new insights into these questions.

Our findings reveal the close relationship between nicotine upregulation of α4β2Rs and dispersal of the GA. By labeling brain sections from the NAc of mice with Abs specific for St3 (Fig. 3D-F), we have validated our *in vitro* findings about nicotine-induced GA dispersal *in vivo*, in VTA dopaminergic neurons where nicotine upregulation of α4β2Rs is well established. Both processes occur simultaneously in α4β2R-expressing HEK cells and cultured neurons, and the dose-dependence of nicotine and the time course of the onset and reversal of these processes are similar (Govind, Walsh et al. 2012). Another relevant set of findings is that specific labeling of intracellular α4β2R high-affinity bindings sites in HEK cells and neurons is only observed in the St3-containing Golgi puncta and is not seen for α4β2Rs in the ER, where they are assembled (Figs. 1G, 2J and supplemental Fig. S2-2A). These findings suggest that the post-translational events that cause the formation of the high-affinity bindings sites occur after α4β2Rs have assembled in the ER and have been trafficked through the secretory pathway and, eventually, through the St3-containing Golgi puncta. In support of this interpretation is our previous finding that nicotine exposure increases the numbers of α4β2R-containing acidic vesicles, and that this perfectly parallels α4β2R upregulation, as assayed by increases in α4β2R high-affinity binding (Govind, Vallejo et al. 2017). This interpretation is further supported by our data showing that as nicotine treatment increases the number of the St3-containing Golgi puncta, it also causes α4β2R upregulation as assayed by changes in the number of α4β2R high-affinity binding sites (supplemental Fig. S5-1B).

The picture that emerges from our findings is that dispersal of the GA by nicotine precedes, and is required for, α4β2R upregulation, and results in a change in the secretory pathway taken by the receptors after release from the ER. After nicotine exposure, α4β2Rs traffic through the altered GA, specifically through the newly formed, small St3-containing Golgi membranes. The formation of high-affinity binding sites on α4β2Rs occurs during trafficking of α4β2Rs through this altered pathway. In the absence of nicotine exposure, newly assembled α4β2Rs are released from the ER, largely traffic through the compact Golgi, and do not form high-affinity binding sites. A small number of St3-containing Golgi puncta do exist in untreated cells (Fig. 1D), and trafficking through the limited numbers of these puncta leads to a small number of high-affinity binding sites in the absence of nicotine exposure. In cultured neurons, there are much higher levels of St3-containing Golgi puncta, especially in dendrites and axons, which are probably formed in response to the synaptic activity-driven dispersal of the GA (Fig. 4). Their presence explains why in the absence of nicotine exposure there are relatively high levels of α4β2R high-affinity binding sites in neurons *in vitro* and *in vivo* (Govind, Vezina et al. 2009, Lewis and Picciotto 2013, Melroy-Greif, Stitzel et al. 2016). The high initial levels of binding sites in neurons also explains why the fold-increase in high-affinity binding sites with nicotine exposure is relatively low compared to that in heterologous expression systems such as HEK cells (Govind, Walsh et al. 2012).

### Is “fragmented” GA in neurons a second functional GA state?

The Golgi apparatus (GA) is often described as a stacked set of disk-like membranes located next to the nucleus at the microtubule organizing center (MTOC). There it functions in the forward trafficking of lipids and proteins in the secretory pathway between the endoplasmic reticulum (ER) and the plasma membrane, or other organelles in the cell. Because several neurodegenerative diseases feature neurons with fragmented GAs, this fragmentation is often assumed to be pathological and deleterious. However, a dispersed or fragmented GA morphology has been shown to be perfectly functional in many cell types, including yeast (Suda and Nakano 2012), plant cells (Vildanova, Wang et al. 2014) and skeletal muscle fibers (Ralston, Lu et al. 1999). The organization of the Golgi complex and microtubules in skeletal muscle is fiber type-dependent (Ralston, Lu et al. 1999) and Golgi membranes are typically highly dispersed in a manner similar to what we have observed in α4β2R-expressing HEK cells after nicotine exposure (Fig. 1). Even in cells with compact, peri-nuclear GA morphology, it progressively changes into a dispersed morphology during mitosis, and reverses after (Ayala and Colanzi 2017). In all of these cases, the dispersed Golgi functions fully as part of the secretory pathway, in a manner similar to intact, perinuclear GA. As others have proposed (Makhoul, Gosavi et al. 2018, Kulkarni-Gosavi, Makhoul et al. 2019) dispersed GA can function as a separate Golgi state, and a dispersed morphology appears to have functional advantages in the cell types where it is found.

Other cells such as neurons (Horton, Yi et al. 2006) and glia (Fu, McAlear et al. 2019) have both a compact peri-nuclear GA in their somata, as well as dispersed Golgi membranes in both their somata and extended processes. The mixed GA morphologies in these cells suggest that the degree to which the GA is found in the dispersed state, relative to the compact state, can vary. Here, we have linked changes in GA morphology in neurons, from a compact to a dispersed or fragmented state, with increases in small, mobile Golgi vesicles throughout dendrites and axons. The reason this phenomenon has gone largely unnoticed is because the smaller Golgi vesicles lack proteins typically used as standard Golgi markers. Using a variety of Golgi glycan-modifying enzymes as markers (Table I), we were able to identify large numbers of additional Golgi membranes in dendrites and axons. Many previous studies have also observed membranes labeled by Golgi glycan-modifying enzymes. However, these structures were thought to have “ectopic” staining of Golgi glycosyltransferase markers (Berger 2002) because they were not labeled by “standard” Golgi markers such as GM130 and giantin.

Our initial observation that long-term nicotine exposure can totally disperse the GA in HEK cells when α4β2Rs were stably expressed (Fig. 1) was advantageous. This discovery provided us with a simple model system for characterizing how a physiological, extracellular stimulation of cultured cells could activate a switch in the state of the GA from a perinuclear compact set of stacks to a dispersed set of membranes widely distributed throughout the cells. It also allowed us to identify similar changes in neurons, where changes in GA morphology were less evident. One significant difference between α4β2R-expressing HEK cells and neurons is that neurons show varying degrees of GA dispersal, particularly in dendrites and axons, and this was most evident when visualized with the Golgi marker, St3. Another major difference between Golgi membranes of α4β2R-expressing HEK cells and neurons is that dendritic Golgi membranes are largely not labeled by standard Golgi markers, such as GM130 (Fig. 2 A and B), and axonal Golgi membranes are not labeled by GM130 at all (Fig. 3A and B). The lack of GM130 staining in dendrites and axons led previous studies to conclude that functional Golgi membranes were largely not present in these compartments (Hanus and Ehlers 2016), and also prevented the observation that different stimuli can regulate the levels and distribution of these membranes. For instance, Thayer et al. (Thayer, Jan et al. 2013) were able to measure dispersal of the somatic GA, as identified by GM130 staining, in response to changes in electrical activity, but did not observe the changes we characterized in dendrites and axons through the use of different Golgi markers.

In neurons, we assume that dispersed Golgi membranes arise after nicotine exposure or increased synaptic activity through forward trafficking out of ERESs, similar to what we observed in the HEK cells (Fig. 7C). Neurons differ from the HEK cells with Golgi being found in dendrites and axons. The Golgi in these processes differ from the somata because membranes lack GM130 staining, except for occasional GM130-containing Golgi outposts in dendrites. Dendrites differ from axons in that Golgi can emerge from ERESs found throughout the dendritic ER, while axons lack ERESs in the ER past the axon initial segment. We found that soluble ER cargo in the soma readily entered and extended through the ER in dendrites, but not in axons, indicating that a diffusional barrier may exist at the axon initial segment. What causes this barrier is not clear, but the narrowing of ER in axons (Terasaki 2018) could contribute.

**Figure 7.**
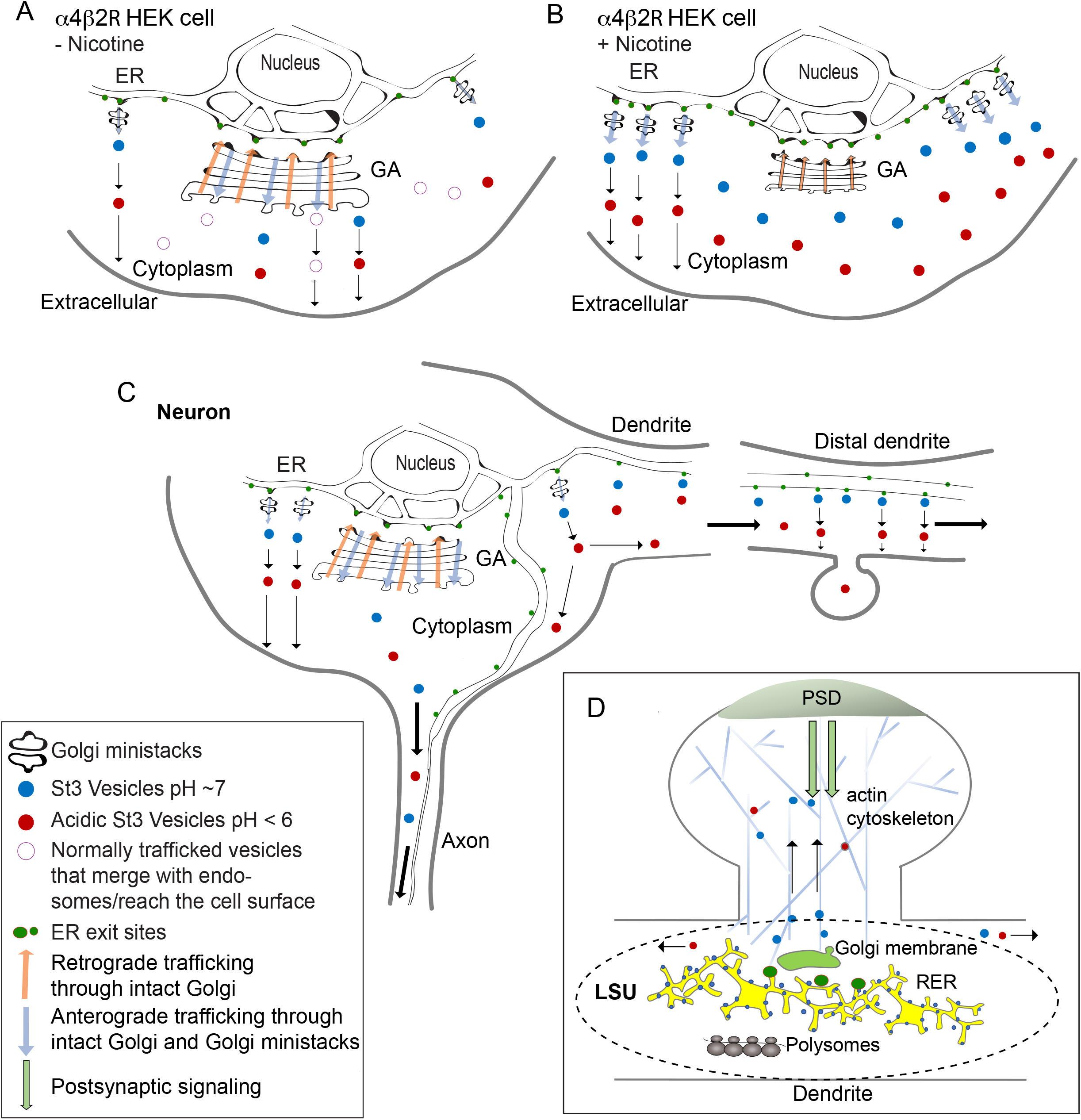
A. Model depicting the change in the GA for α4β2R HEK cells under untreated conditions (-Nicotine; top). B. Golgi dispersal and increased mini Golgi vesicles arising from ER exit sites in nicotine treated α4β2R HEK cells (+ Nicotine; bottom). C. Schematic representation of GA and its distribution in cultured neurons. D. Model of local secretory unit (LSU) and its potential role in synaptic tagging and capture.

### Extracellular signaling regulates the state of the GA

Overall, our results suggest that different types of physiological, extracellular signals can trigger signal transduction mechanisms that control transitions between the different GA states. The ability of extracellular stimuli, such as nicotine exposure, to regulate the state of the GA is a central feature of the model displayed in Fig. 7A and B for α4β2R-expressing HEK cells, and Fig. 7C for neurons. Previous supporting evidence includes the physiologically-induced changes in Golgi morphology and distribution that occur with different types of cell polarization (Yadav, Puri et al. 2009), such as during fibroblast migration (Yadav and Linstedt 2011), wound healing, and antigen presentation to T cells (Ravichandran, Goud et al. 2020). In these examples, repositioning of the GA directs the secretory pathway so that secretion is largely targeted to a limited plasma membrane domain. This helps to establish and then maintain specialized membrane domains critical for polarization events in response to extracellular signaling. To accomplish this, the GA responds to incoming signaling from the plasma membrane, resulting from cues from the extracellular environment.

The mechanisms underlying nocodazole dispersal of the GA are well-characterized (Cole, Sciaky et al. 1996, Storrie, White et al. 1998, Fourriere, Divoux et al. 2016). To begin to characterize the mechanisms causing nicotine-induced GA dispersal in HEK cells and neurons, we performed a detailed comparison of the GA dispersal induced by either drug (Fig. 5). The α4β2R-expressing HEK cell system was used to characterize in more detail how nicotine treatment disperses the GA. We found that nicotine and nocodazole dispersal are very similar in terms of the size and Golgi marker distribution of the dispersed Golgi membranes (Fig. 5B; Supplemental Fig. S5-1A) and their alignment with ERESs (Supplemental Fig. S5-3A). These findings, together with the demonstration that cargo moves directly from the ER through the dispersed Golgi membranes in HEK cells (Fig. 5D) and in dendrites (Fig. 6), support the models shown in Fig. 7.

Our model assumes that the initial steps of GA dispersal in response to nicotine are the same as those for nocodazole-induced dispersal. In the absence of nicotine exposure, the GA in non-dividing HEK cells is in the compact perinuclear state (Fig. 7A). Forward trafficking of cargo from the ER largely involves movement out of ERESs situated just apposed to the cis-Golgi stack. At steady-state, the forward trafficking into the Golgi is countered by exit from the trans-Golgi to the plasma membrane and other organelles, along with retrograde trafficking from the cis-Golgi back to the ER (Cole, Sciaky et al. 1996). In our model, nicotine exposure alters the steady-state balance, as forward trafficking out of the former ERESs becomes highly reduced (Fig. 7B). After GA dispersal, cargo emerges from more peripheral ERES, initially forming ministacks instead of trafficking to the compact GA and fusing to form the cis-most stack. The ministacks go through a process of polarization, after which cargo undergoes normal

GA processing and is then trafficked elsewhere (Fourriere, Divoux et al. 2016). We have identified some of these events via imaging of live cells (Fig. 1E), which demonstrated that St3-containing mobile vesicles exit from the trans side of polarized mini-stacks following nicotine-induced GA dispersal. We have also observed that only a subset of St3-containing mobile vesicles have acidic lumens, as indicated by pHrodo staining (Fig 1F). This observation suggests that after St3-containing vesicles exit the ministack, they undergo further maturation, during which their lumen becomes acidified. These maturation events after are depicted in the model as a transition between the less mature St3-containing vesicles (blue) and the acidic St3-containing vesicles (red). We propose that it is the acidic St3-containing vesicles that traffic to the plasma membrane.

In our model (Fig. 7B), dispersed Golgi membranes arise after nicotine exposure through forward trafficking out of ERESs, not from uncoupling and/or fission of the GA stack, as has been proposed for the fragmentation associated with different neurodegenerative diseases (Machamer 2015, Makhoul, Gosavi et al. 2019, Martinez-Menarguez, Tomas et al. 2019) and the formation of Golgi outposts (Quassollo, Wojnacki et al. 2015). Unlike nocodazole-induced GA dispersal, the downstream effects of nicotine do not involve depolymerization of the microtubule network in either HEK cells (Fig. 5C) or cultured neurons (Fig. 5 E and G). Nicotine-regulated signaling pathways may target other microtubule-related processes, such as the activity or attachment of the microtubule motor, dynein, which traffics cargo from ERESs to the cis-stack of the compact perinuclear GA. Blocking of dynein-mediated transport has been found to disperse the GA (Fourriere, Divoux et al. 2016). In the model (Fig. 7A), we suggest that the dispersed GA tied to more peripheral ERESs may normally exist at low levels in cells and contribute to forward trafficking of secreted lipids and proteins. Consistent with low levels of dispersed Golgi in the absence of nicotine exposure is the observation that high-affinity binding sites form on α4β2Rs in the absence of nicotine exposure. As discussed above, high-affinity binding sites appear to form on α4β2Rs only when they traffic through dispersed Golgi.

### Dispersed Golgi membranes in dendritic local secretory units and synaptic tagging

Because of their complex morphology, the polarization of pyramidal neurons during development occurs in stages, as demonstrated both *in vitro* (Dotti, Sullivan et al. 1988) and *in vivo* (Calderon de Anda, Gartner et al. 2008). In parallel with this process, the GA in neurons undergoes a series of changes in positioning and morphology as different regions of the neuron polarize and develop. First, during specification of the axon, the GA and MTOC reposition so that the GA is oriented closely to the axon initial segment. Then, as dendrites develop, the Golgi is again repositioned so that it is placed at the initial segment of the leading dendrite, often extending into the dendrite (Horton, Racz et al. 2005). The appearance of Golgi outposts (defined by the presence of the Golgi marker, GM130) occurs after dendrite maturation when synaptogenesis is occurring. The formation of synapses begins the process of neurotransmitter release onto postsynaptic sites and results in increased excitatory synaptic input to the dendrites. We have demonstrated that increased excitatory synaptic input leads to increased levels of small Golgi membranes in dendrites (Fig. 4). These Golgi membranes could function at local secretory sites, where, together with ribosomes, they can mediate local translation of secreted and membrane proteins, as indicated in Fig 6. During synaptogenesis, we would expect the initiation and ramping up of synaptic activity to recruit Golgi elements, ribosomes and ER membranes required for the assembly of local secretory units (LSUs). Similar to other examples of GA repositioning during neuronal development, the Golgi component of the LSU would facilitate the local membrane polarization needed for synapse maturation and maintenance at developing postsynaptic sites.

A central question in neuroscience is how changes in postsynaptic activity in dendrites are integrated with other neural activity and result in functional plasticity over the short-term, and structural plasticity over the long-term. The process of protein translation and its local control has long been implicated in long-term structural plasticity changes (Smolen, Baxter et al. 2019). Our findings of previously undetected functional Golgi membranes throughout dendrites raises new possibilities of the existence of large numbers of LSUs that service postsynaptic sites. In addition, the ability of synaptic activity and nicotine exposure to increase Golgi levels and their distribution in dendrites suggests that the numbers and placement of LSUs are regulated by local signaling at synapses. The Golgi membranes in LSUs would play a critical role in a number of LSU functions. First, Golgi would be needed for the local control of forward trafficking of secreted and membrane proteins specifically targeted to synapses. This would include proteins locally synthesized from transported mRNAs and proteins, such as NMDA receptors, and those that are synthesized in the soma and specifically trafficked to local Golgi membranes at synapses (Jeyifous, Waites et al. 2009). Second, Golgi membranes serve as local organizers of the actin and microtubule cytoskeleton (Ravichandran, Goud et al. 2020), which would be critical for the placement and regulation of vesicle trafficking into and out of the postsynaptic membrane. Third, Golgi membranes can act as local signaling scaffolds for the integration of signaling from local domains (Makhoul, Gosavi et al. 2018, Kulkarni-Gosavi, Makhoul et al. 2019).

Another potential function of LSUs at postsynaptic sites is to serve as coordinating sites in the process of synaptic tagging. For many years, the concept of synaptic tagging and capture has emerged from studies of long-term memory consolidation and the late phase of long-term potentiation at synapses, which is dependent on protein synthesis (Davis and Squire 1984, Smolen, Baxter et al. 2019). Early models of the molecular events underlying synaptic tagging proposed a soluble tag molecule in the cytoplasm of dendrites that would bind to scaffolded receptors inserted into the postsynaptic domain in order initiate the tagging of that synapse (Redondo and Morris 2011). This concept of a yet-to-be discovered soluble synaptic tag and its receptor continues today in synaptic tagging models, despite a lack of any direct evidence for either. An alternative model for how synaptic tagging and capture could be achieved is schematized in Fig. 7D. We propose that a more realistic mechanism is an LSU strategically placed in the dendritic shaft to coordinate the incoming trafficking of mRNA particles, ribosomes, cytoskeletal elements, trafficking vesicles, and the placement of mitochondria, ER and degradative machinery. LSU strategic placement would depend on incoming local synaptic signaling and trafficking of the various LSU components from the soma. The LSU would coordinate all of these elements among a variable number of postsynaptic membranes in the vicinity and set the distribution based on local synaptic weights. The synaptic weights would be set based on individual inputs from each postsynaptic domain, as well as homeostatic input arriving from the soma. All of these inputs would be integrated at the LSU, allowing constant readjustment of the distribution to each domain. The Golgi element would play a central role through its function in the forward trafficking of secreted and membrane proteins, organization of the local cytoskeleton to each domain, and its scaffolding of intracellular signaling molecules.

## Materials and Methods

### Antibodies, cDNA constructs and reagents

Antibodies used were either purchased from commercial suppliers or were generous gifts from various researchers. Antibodies against the following antigens were used, GM130 (monoclonal; BD transduction laboratories, CA), GM130 (polyclonal; Sigma, MO), HA epitope (HA.11, monoclonal, Biolegend,CA), St3 Gal3 (polyclonal; Abcam, MA), Tyrosine Hydroxylase (BD transduction laboratories, CA), NFH (EMD Millipore, MA) and St3 Gal5 (polyclonal; CS14 Smith affinity purified) gift from Michael Tiemeyer (University of Georgia). The Fluorescent secondary antibodies and ligands that were used include Alexa Fluor® 488-conjugated goat anti-rabbit IgG, Alexa Fluor® 568-conjugated goat anti-mouse IgG, Alexa Fluor® 568-conjugated goat anti-rabbit IgG, Alexa Fluor® 647-conjugated goat anti-mouse IgG, Alexa Fluor® 647-conjugated goat anti-rabbit IgG (Thermofisher scientific, MA) and Janelia Fluor 549® Halo ligand (Promega, WI). Rat α4 and β2 used for generating the stable cell line were provided by Dr. Jim Boulter (University of California, Los Angeles, CA). The HA epitope, YPYDVPDYA, and a stop codon were inserted after the last codon of the 3′-translated region of the subunit DNA of the β2 using the extension overlap method as described in Vallejo *et al*. (Vallejo, Buisson et al. 2005). Other cDNAs generously provided include: St3-eGFP from Dr. Sakari Kellokumpu, (University of Oulu, Oulu, Finland), GalNAc T2-mCherry from Dr. Dibbyendu Bhattacharya (Tata Memorial Center, Mumbai, India), GalT-3xGFP and mCherry-Sec23 from Dr. Jennifer Lippincott-Schwartz (HHMI, Janelia Research Campus), GM130 GFP from Dr. Christine Suetterlin (University of California, Irvine, CA), pDsRed-ER (Takara Bio, CA), RUSH-GPI-mApple, Rush-GPI-Halo, and Ensconsin-mCherry (Addgene, MA). For imaging ER bulk flow trafficking to Golgi, a bicistronic plasmid encoding a modified DsRed-Express2 cargo (Casler, Papanikou et al. 2019) and ManII-GFP was generated in the laboratory of Benjamin Glick (University of Chicago, Chicago, IL). The M. musculus ManIIA1gene was obtained from BioBasic (Toronto, Canada), PCR amplified and inserted into the DsRed-Express2 plasmid using in-fusion cloning (Takara Bio, CA). St3-Halo was generated by replacing the GFP in St3-eGFP with Halo tag (GenScript USA Inc, NJ). The following reagents were used: Nifro Rhodamine (developed in the laboratory of Dr. Jogesh Mukherjee (University of California, Irvine, CA), pHrodo (Thermofisher scientific, MA), (-)-Nicotine, Nocodazole, Bicuculline (Sigma-Aldrich, MO), DL-APV (Abcam, MA) and ProLong™ Gold Antifade Mountant with DAPI (Thermofisher scientific, MA).

### Mammalian cell culture and transfection

The human embryonic kidney (HEK 293T) cell line stably expressing the large T antigen (tSA201 cells) was from Dr. J. Kyle (University of Chicago, Chicago, IL). This cell line is not in the list of Database of Cross-Contaminated or Misidentified Cell Lines. Using this parent HEK 293 T cells, a stable cell line expressing rat α4β2 nAChRs was generated in our lab, expressing untagged α4 and C-terminal, HA epitope-tagged β2 subunits (Vallejo, Buisson et al. 2005). Both the parent HEK and stable α4β2 HEK cell lines were maintained in DMEM (Gibco, Life technologies) with 10% calf serum (Hyclone, GE Healthcare Life Sciences, UT) at 37°C in the presence of 5% CO_2_. DMEM was supplemented with Hygromycin (Thermofisher scientific, MA) at 0.4 mg/ml for maintaining selection of α4β2 HEK cells. Hoechst staining and immunofluorescent detection were performed periodically to test for mycoplasma contamination. Fresh batches of cells were thawed and maintained only up to two months.

HEK cells were grown on 22X22 mm coverslips (Thermofisher scientific, MA) or 35 mm imaging plates (MaTek, MA) coated with poly-D-lysine. Stable cells were plated in media without hygromycin for experiments. 75% confluent cultures were transfected with 0.5µg DNA of indicated constructs with Lipofectamine 2000 transfection reagent (Thermofisher scientific, MA). After 24 hours of transfection cells were treated with 10 µM nicotine for 17 hours.

### Primary neuronal culture and transfections

Primary cultures of rat cortical neurons were prepared as described (Govind, Walsh et al. 2012) using Neurobasal Media (NBM), 2% (v/v) B27, and 2 mM L-glutamine (all from Thermofisher Scientific, MA). Dissociated cortical cells from E18 Sprague Dawley rat pups were plated on slips or plates coated with poly-D-lysine (Sigma, MO). For live imaging, neurons were plated in 35 mm glass bottom dishes (MatTek, MA). Cells were plated at a density of 0.25X10^6^ cells/mL on 35 mm dishes or per well in a 6-well plate. Neuronal cultures were transfected at DIV 10 with the Lipofectamine 2000 transfection reagent (Thermofisher Scientific, MA) according to manufacturer’s recommendations. Neurons were transfected with cDNAs of α4, β2_HA_ and various Golgi markers, ER/ER exit site markers or RUSH constructs. 0.5 ug of each DNA up to a total of 2ug were used per 35 mm imaging dish or per well of a 6 well plate. 24 hours after transfection, neurons were treated with 1 μM nicotine for 17 hrs. Bicuculline and DL-APV treatments were performed as described previously (Thayer, PNAS, 2013).

### Immunocytochemistry

HEK cells or neurons grown in imaging chambers (MaTek) were live imaged in low fluorescence Hibernate E buffer (Brain Bits, IL). Those grown on coverslips were fixed with 4% paraformaldehyde + 4% sucrose for 10 minutes. Surface expressed α4β2 receptors were imaged by live labeling the cells with mouse anti-HA antibody (1:500) for 40 minutes prior to fixation. Cells were permeabilized with 0.1% triton X-100. Primary antibody incubations were for 1 hour and secondary antibodies for 45 minutes unless otherwise mentioned. Coverslips were mounted using Prolong Gold mounting media with DAPI.

Fluorescence images were acquired using a Leica SP5 Tandem Scanner Spectral 2-Photon scanning confocal microscope (Leica Microsystems), or Marianas Yokogawa type spinning disk confocal microscope with back-thinned EMCCD camera. Images were processed and analyzed using ImageJ/Fiji (US National Institutes of Health).

### *In vivo* nicotine experiments

Mice were housed in standard conditions on a 12:12-h light-dark cycle in a temperature- and humidity-controlled facility and allowed ad libitum access to standard chow and water. All procedures were in accordance with the guidelines of, and approved by, the Institutional Animal Care and Use Committee at the University of Chicago. Mice received either 100 µg/ml nicotine via drinking water, or regular water, daily for 3 weeks. Nicotine intake was monitored daily and treatment did not alter water intake or body weight. Nicotine doses were chosen based on previous research showing that when administered via drinking water, the nicotine concentrations correlated with nicotine and cotinine blood levels that induce nicotine tolerance and dependence in mice (Koranda, Cone et al. 2014).

### Immunohistochemistry

Immunostaining was performed as described previously (Koranda, Krok et al. 2016). Briefly, brain tissues were fixed with 4% formaldehyde overnight and transferred into 30% sucrose for 24 hrs. 40 μm coronal serial brain sections were made using a cryostat (Leica Instruments). Tissues underwent heat-mediated epitope retrieval for 30 min with a buffer composition of 10 μM sodium citrate, pH 9 in Tris-buffered saline (TBS). After blocking in 0.05 M Tris-buffered saline containing 5% normal donkey serum and 0.3% Triton X-100 for 1 h at room temperature, sections were transferred to primary antibody, rabbit St3 pAb (1:300; Abcam) and mouse TH (1:750), containing 0.3% Triton X-100 with 3% BSA and incubated at 4°C for 48 h. The secondary antibody, anti-rabbit Cy3 (1:500; Jackson ImmunoResearch), was diluted in 3% BSA and sections were incubated for 1 h at room temperature.

Imaging was performed using the SP5 Leica confocal system, with a 63x oil objective lens with a resolution of 0.074um x 0.074um x 0.130 µm. After acquisition, images were deconvolved using Huygens Professional software using classic maximum likelihood estimations (Scientific Volume Imaging). Imaging and analysis were performed by an analyst blind to the experimental conditions.

### Image Analysis

St3 punctal densities were quantified using maximum *z* projections of background-subtracted images. Thresholded punctal size and density was analyzed using the Analyze Particle function in Fiji/ImageJ. Results are expressed as mean ± SEM of *n* samples unless stated otherwise. Statistical comparisons were made using two-tailed Student’s *t* tests or ANOVA/Tukey *post hoc* analysis as indicated. Statistical graphs were generated with StatPlus software.

### Live 3D-SIM imaging

3D-SIM imaging experiments were performed on a custom-built, 3D-SIM microscope system housed in the Advanced Imaged Center (AIC) at the Howard Hughes Medical Institute Janelia research campus. System configuration and operation have been previously described (Li, Shao et al. 2015). Briefly, dual color images (488- and 560-nm laser lines) were acquired using a Zeiss AxioObserver microscope outfitted with a Zeiss 100x, 1.49 NA oil objective and two cameras. System hardware and image reconstruction was achieved using custom software written and provided by the AIC.

## Acknowledgment

This work was financially supported by NIH RO1 DA035430 (W. N. G.), DA044760, (W. N. G. and J. M.), DA043361 (X. Z and W. N. G.) and Peter F. McManus Foundation (W. N. G.). Structured illumination microscopy (SIM) imaging data used in this publication was produced in collaboration with the Advanced Imaging Center (AIC), a facility jointly supported by the Gordon and Betty Moore Foundation and HHMI at HHMI’s Janelia Research Campus. Authors would like to thank Teng-Leong Chew, Satya Khuon and Aaron Taylor at the AIC, Janelia Research Campus for technical support and SIM data acquisition. We thank Vytas Bindokas (University of Chicago) and Louie Kerr (Marine Biological Laboratory), Panagiotis Chandris and Hari Shroff (NIH/NIBIB) for technical support on microscopy; Luke Lavis (Janelia research campus) for Halo dyes and Dibbyendu Bhattacharya (Tata Memorial Center), Sakari Kellokumpu (University of Oulu), Jim Boulter (University of California), Steven Standley (Western University of health sciences) for cDNA constructs used in this study. We thank U of C undergraduates Briana Turner, Abhijit Ramaprasad and William Ramos for assistance. The authors declare that they have no competing interests.

