## Supplemental Data for "Nicotine exposure and neuronal activity regulate Golgi membrane dispersal and distribution"

### Supplementary Figures

Suppl Figure S1-1

A. 3D Structured illumination (3D-SIM) images of $\alpha4\beta2$-expressing HEK cells. Cells were transfected with St3-GFP for 24 hours, treated with or without 10 $\mu$M nicotine for 17 hours and live-imaged. Scale bar, 10 $\mu$m. Inset Scale bars (right), 5 $\mu$m. B. Histograms displaying the size distribution of St3/GM130 (top) and St3-only (bottom) puncta (8-10 cells per group). C. Classification and quantification of cells into three categories (intact, partially dispersed, and fully dispersed) based on Golgi integrity and morphology. $\alpha4\beta2$ HEK cells were transfected with St3-GFP (green) for 24 hours and treated with or without 10 $\mu$M nicotine for 17 hours. Cells were fixed and stained with DAPI (blue). Top, representative images of cells with intact, partially dispersed and fully dispersed Golgi. Bottom, percentage of St3-GFP transfected cells with each phenotype were quantified under nicotine treated and untreated conditions. Data represents $\pm$ SEM (n=4; total number of St3-GFP transfected cells analyzed; control cells, 487; nicotine cells, 469; percentage of cells with intact Golgi, control cells, 69 $\pm$ 1.7%; nicotine cells, 23.3 $\pm$ 3.6%, con vs nic, **$p \leq 0.0013$; partially dispersed Golgi, control cells, 23.3 $\pm$ 1%; nicotine cells, 32.3 $\pm$ 3.4%, con vs nic, ns, $p>0.1$; dispersed Golgi, control cells, 6.2 $\pm$ 2.2%; nicotine cells, 45.2 $\pm$ 3.2%, con vs nic, **$p \leq 0.0019$)

Suppl Videos S1-2; A, B, C

Live-imaging movies of St3 mobility in HEK cells for intact (A), partially dispersed (B), and fully dispersed (C) Golgi phenotypes. Image frames were acquired every sec for 40-60 sec.

Suppl Figure S2-1

A. Variable nature of somatic Golgi morphology in primary cortical cultures. DIV 10 neurons were transfected with St3-GFP (green) for 24 hours and imaged after fixation. Representative images depicting variable distribution of St3 in neurons. Scale bar, 10 $\mu$m.

Suppl Figure S2-2

Distribution of $\alpha4\beta2$ nicotinic receptor in primary cortical cultures. A. DIV 10 neurons were transfected with St3-GFP (green) and nicotinic receptor subunits $\alpha4$ and $\beta2$HA for 24 hours, and treated with 1$\mu$M nicotine for 17 hours. A. Cultures were labeled with 25 $\mu$M NifroRhodamine (red) for 30 minutes and imaged live. B. Cultures were fixed, permeabilized and stained with anti-HA antibody (red; secondary antibody anti-mouse Alexa 568). Scale bar, 10 $\mu$m.

Suppl Figure S2-3

Mobility of St3 in dendrites and axons. DIV11 cultured cortical neurons were transfected with cDNA encoding St3-GFP and subjected to live-imaging. A. An example of mobile St3-GFP puncta in the dendrites of a cultured cortical neuron. Dendritic St3-GFP puncta moved with an average velocity of ~ 1.0 μm/sec (arrowheads indicate mobile puncta). Image frames were acquired every 1.5 sec. B. Axonal St3-GFP puncta were mobile in both directions and moved with an average velocity of ~ 1.0 μm/sec. Frames were acquired every 1.5 sec.

Suppl Videos S2-4 A and B
Live-imaging movies of St3 vesicle mobility in dendrites at low (A) and high (B) magnification. Image frames were acquired every sec for 40 sec (A) or 100 sec (B).

Suppl Figure S3-1
Axonal distribution of Golgi marker enzymes. A. DIV11 cultured cortical neurons were either labeled with St3-Gal5 pAb (green, left panels) or transfected with cDNA encoding Man-GFP (green, right panels). Axons were identified by staining with anti-NFH (red). Both endogenous St3-Gal5 and Man-GFP exhibited a punctal distribution in the axons of cultured cortical neurons. The punctal distribution of both markers in distal axonal segments is displayed in the bottom merged panels. Similar to the St3 pAb, the St3-Gal5 pAb does not label somatic Golgi stacks. Scale bars, 10 μm. B. Co-localization of St3 with St3-Gal5 (left) and Man-GFP (right) in axons. Cultures were transfected with cDNA encoding St3-Halo alone (left) or with Man-GFP (right), and stained with St3-Gal5 pAb (left), then visualized to determine overlap of markers. St3-Gal5 and Man-GFP both displayed extensive punctal overlap with St3-Halo in axons (arrows). Scale bars, 5 μm. C. Overlap of St3 and β2 subunits in axons of primary cortical cultures. Cultures were transfected with cDNA encoding St3-GFP and α4β2HA for 24 hrs, then fixed and stained with Abs against HA and NFH. Co-localizing puncta are indicated by arrows. Scale bar, 5 μm.

Suppl Videos S3-2 A and B
Two live-imaging movies of St3-Halo vesicle mobility in axons. DIV11 cultured cortical neurons were transfected with cDNA encoding St3-Halo, labeled with HaloTag ligand, and live-imaged. Image frames were acquired every sec for 1 min.

Suppl Figure S5-1
A. Histograms displaying the size distribution of St3/GM130 (top) and St3-only (bottom) puncta (8-10 cells per group). α4β2 HEK cells were treated with 10 μM nicotine for 17 hours, and further treated for 4 hours with 25 μM nocodazole. Parallel sets of cells were left untreated, treated with nicotine alone (17 hours), or nocodazole alone (4 hours). B. α4β2 HEK cells were treated with nicotine and nocodazole as in A. Cells were washed and bound with 5 nM $^{125}$I epibatidine for 20 minutes, then harvested onto filter paper. Bound radioactivity was measured using a gamma counter. Samples bound with radioactivity in the presence of 1 mM nicotine served to subtract background. Data represents $\pm$ SEM fold change over untreated control, nicotine cells, 6.68 $\pm$ 0.5; nocodazole cells, 1.12 $\pm$ 0.13; nicotine+nocodazole cells, 7.8 $\pm$ 0.56 (n=4; con vs nic, ***$p<0.00002$; con vs noc, ns; con vs nic + noc, ***$p<0.000008$; nic vs nic + noc, **$p<0.008$).

Suppl Figure S5-2
A. Time course of nicotine-dependent Golgi dispersal. α4β2 HEK cells were transfected with Galactosyl transferase-GFP (GalT; green) for 24 hours and exposed to 10 μM

nicotine for the indicated time. Cells were fixed and stained with DAPI (blue). B. Live-imaging of the onset (top panels) and reversal (bottom panels) of nicotine-induced Golgi dispersal. α4β2 HEK cells were transfected with St3-GFP for 24 hours prior to imaging. For onset, 10 μM nicotine was added at time 0 (0h). For measuring reversal, cells were initially exposed to nicotine for 4 hours to induce Golgi dispersal, then nicotine was washed out and imaging performed in drug-free buffer for the indicated time. C. Quantification of the reversal of nicotine-induced Golgi dispersal. α4β2 HEK cells were transfected with St3-GFP (green) for 24 hours and treated with or without 10 μM nicotine for 1 day. To measure reversal, cells were washed with media to remove nicotine and maintained in drug-free media for two days. A parallel set of nicotine-treated cells were maintained in nicotine for the duration of the experiment. Untreated cells were maintained in nicotine-free media. Representative images (top) and quantification (bottom) of the cells under three conditions. Cells were classified and quantified into one of three categories (intact, partially dispersed, and fully dispersed) based on Golgi integrity and morphology (calculated from 10 fields, and 12-60 total cells per treatment group).

Suppl Figure S5-3
A. 3D Structured illumination (3D-SIM) images of α4β2 HEK cells transfected with St3-GFP and ER marker, DsRed-ER (left) or ER exit site marker, mCherry-tagged Sec 23 (mCh-Sec 23; right). Scale bar, 10 μm. Inset scale bar, 2.5 μm (bottom). B. Distribution of DsRed-ER (top) or mCh-Sec 23 (bottom) with St3-GFP in the somata of cultured neurons. C. ERESs, denoted by Sec23, are closely apposed to St3-GFP puncta in the dendrites. Scale bar 10 μm. Inset scale bar 2.5 μm. D. Distribution of ERES marker, Sec 23, in axons and dendrites of primary cortical cultures. DIV 10 neurons were transfected with St3-GFP and mCh-Sec23 for 24 hours. Large arrows show St3-GFP distribution throughout the axon. Small arrows indicate the restricted distribution of Sec23 in the initial segment of axon.

Suppl Figure S6
Secretory trafficking of cargo through dispersed dendritic Golgi membranes. Live-imaging of dendrites showing transport of GPI-RUSH-Halo to plasma membrane occurring via dispersed Golgi, following biotin-mediated release from ER. DIV 10 primary cortical cultures were transfected with St3-GFP and GPI-RUSH-Halo for 24 hours. After biotin addition, GPI-RUSH-Halo entered and accumulated in dispersed, static Golgi structures (St3-GFP), then exited and trafficked to the dendritic plasma membrane. Image frames were acquired every 3 mins for 0.5 h, scale bar, 5 μm.

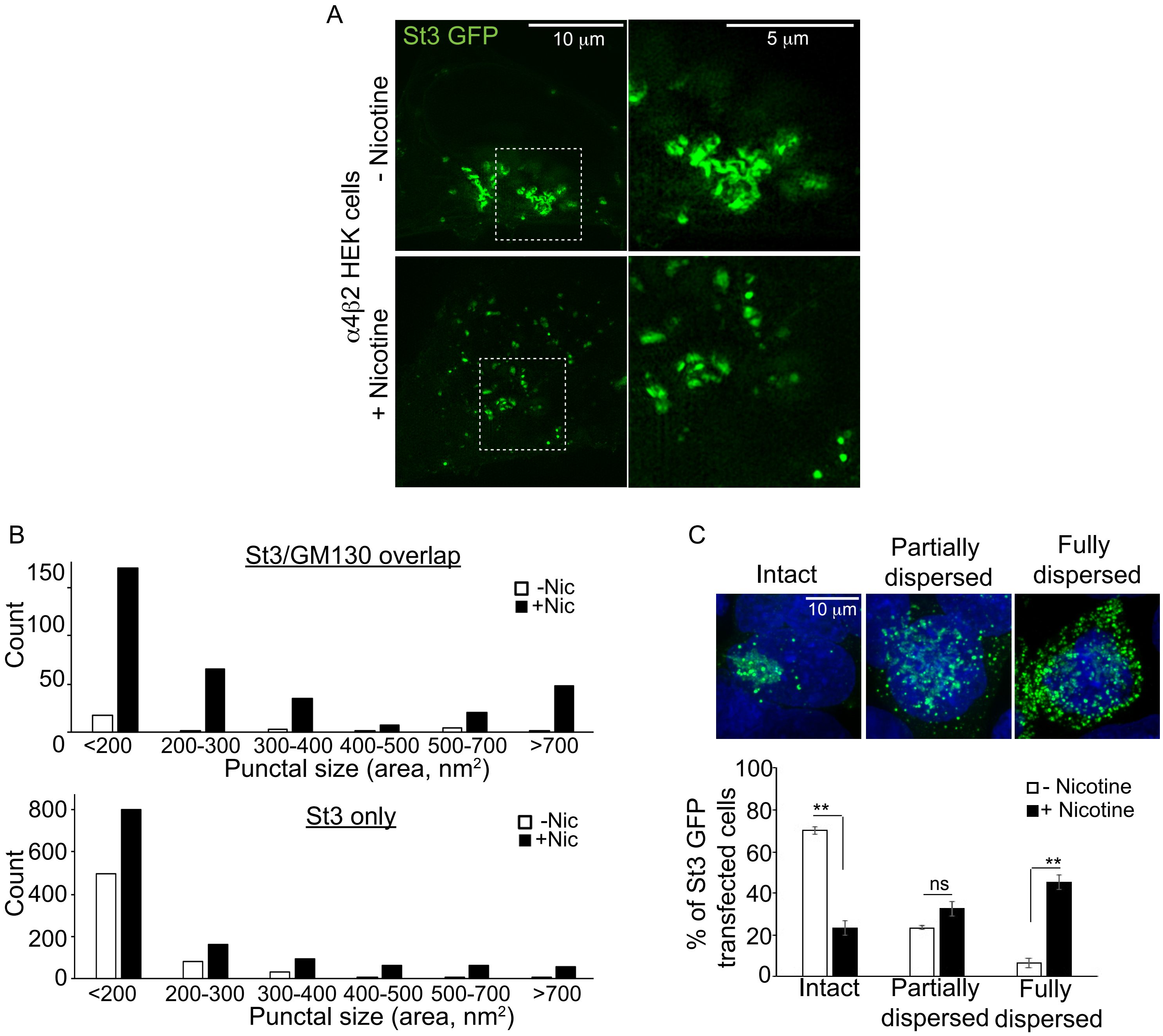
A
St3 GFP
10 μm
5 μm
α4β2 HEK cells
- Nicotine
+ Nicotine
B
St3/GM130 overlap
Count
-Nic
+Nic
<200
200-300
300-400
400-500
500-700
>700
Punctal size (area, nm²)
St3 only
C
Intact
Partially dispersed
Fully dispersed
10 μm
% of St3 GFP transfected cells
- Nicotine
+ Nicotine
**
ns

A

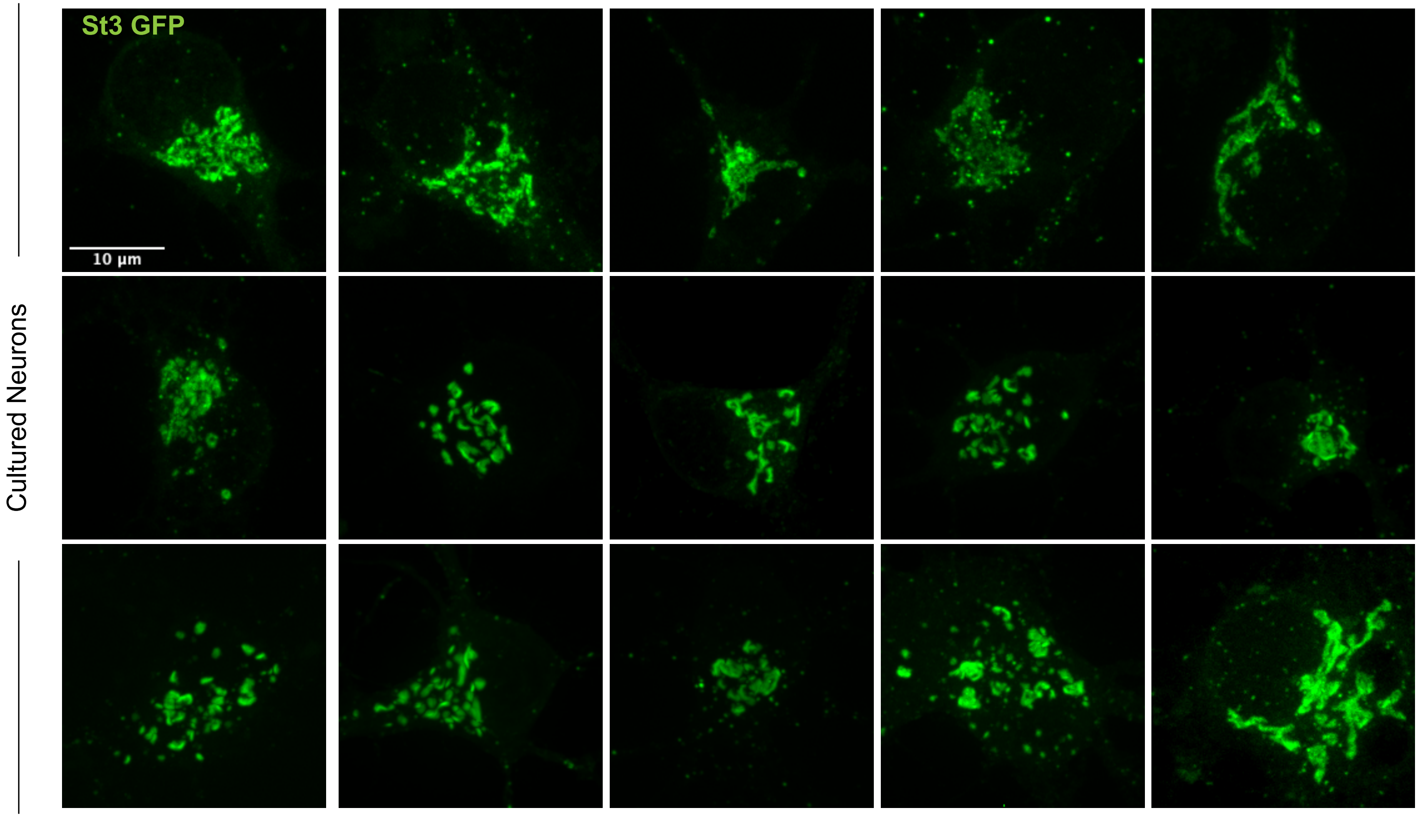

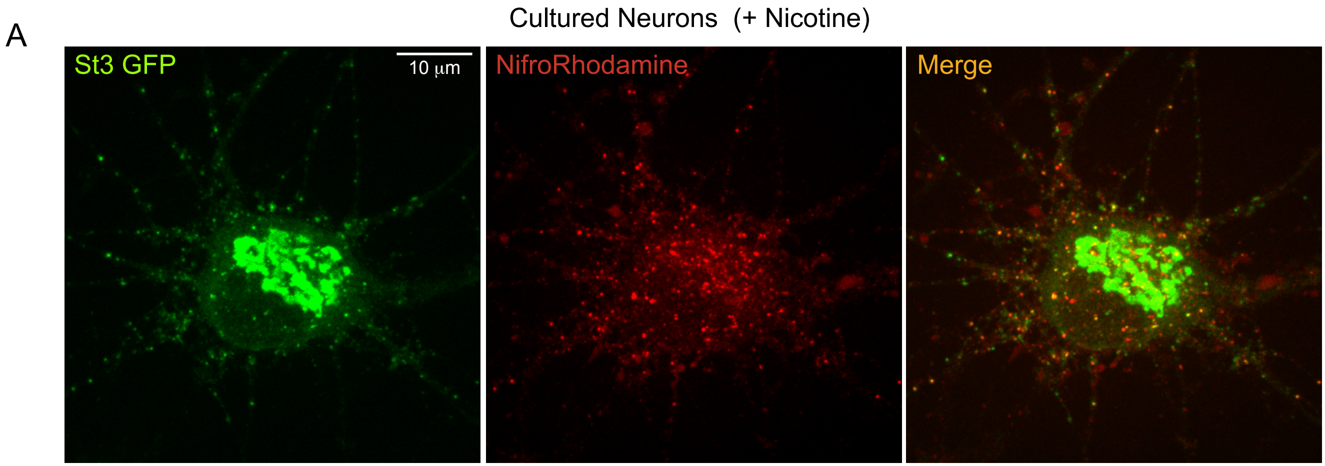
A
Cultured Neurons (+ Nicotine)
St3 GFP
10 µm
NifroRhodamine
Merge

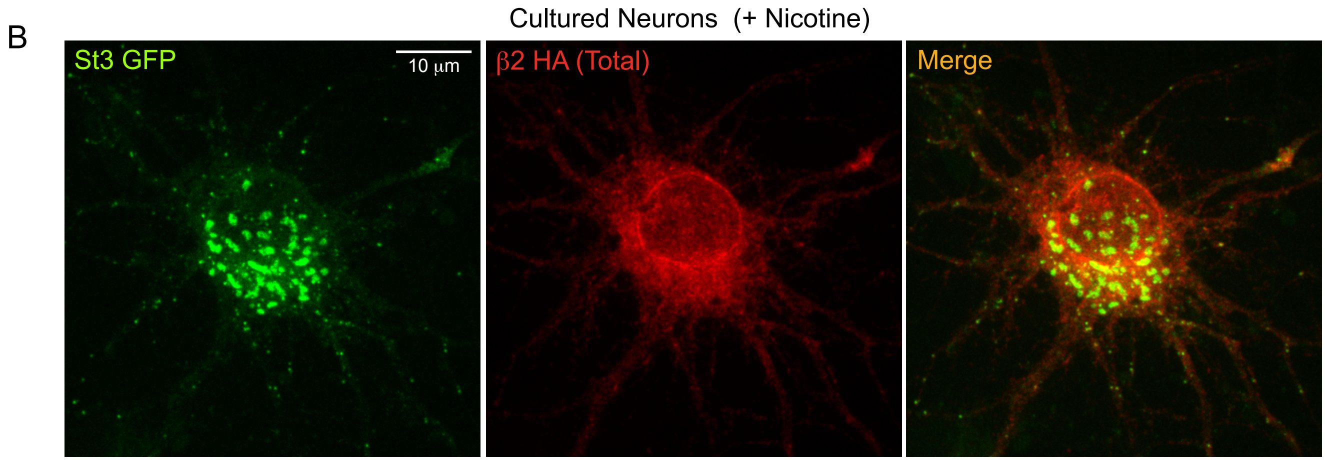
B
Cultured Neurons (+ Nicotine)
St3 GFP
10 µm
β2 HA (Total)
Merge

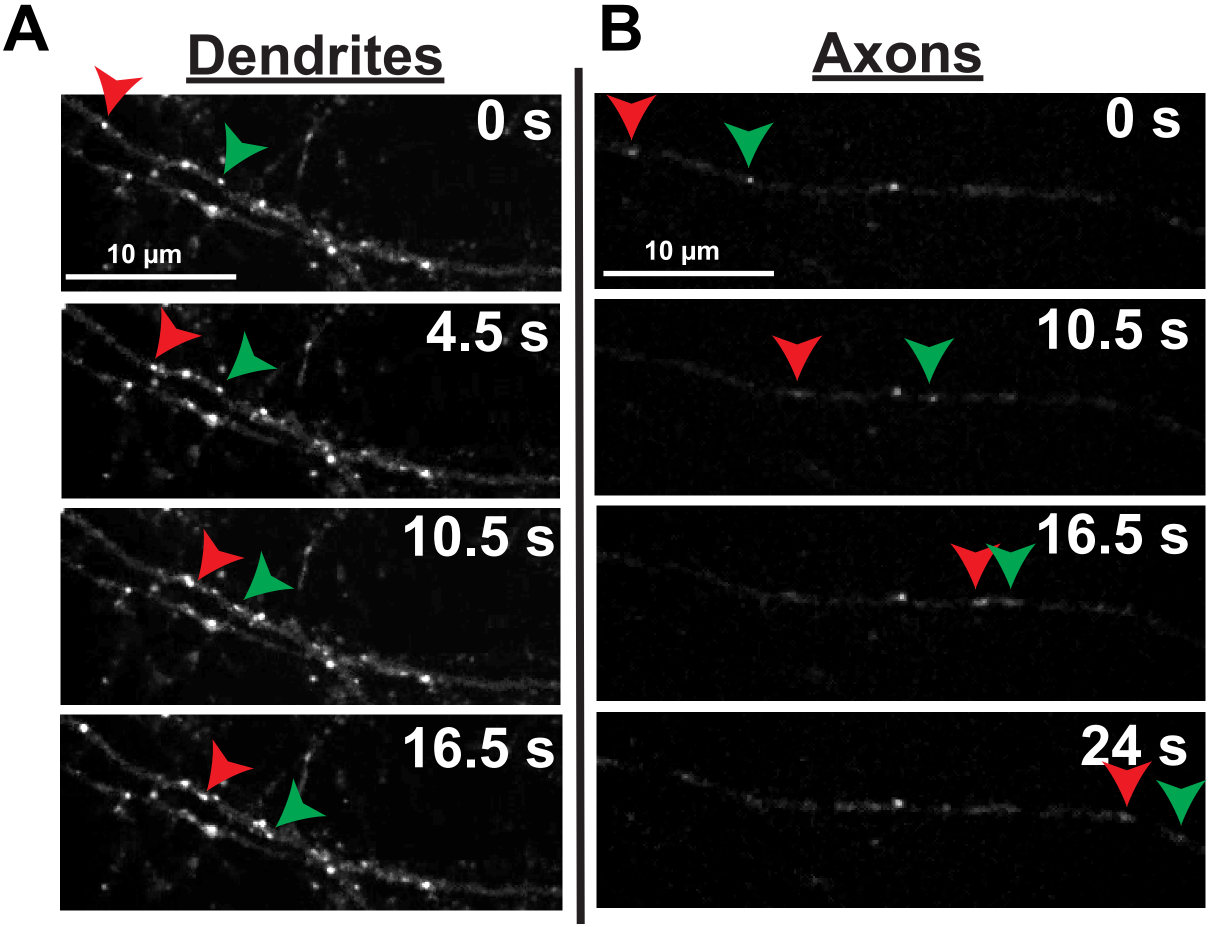
A
Dendrites
0 s
10 µm
4.5 s
10.5 s
16.5 s
B
Axons
0 s
10 µm
10.5 s
16.5 s
24 s

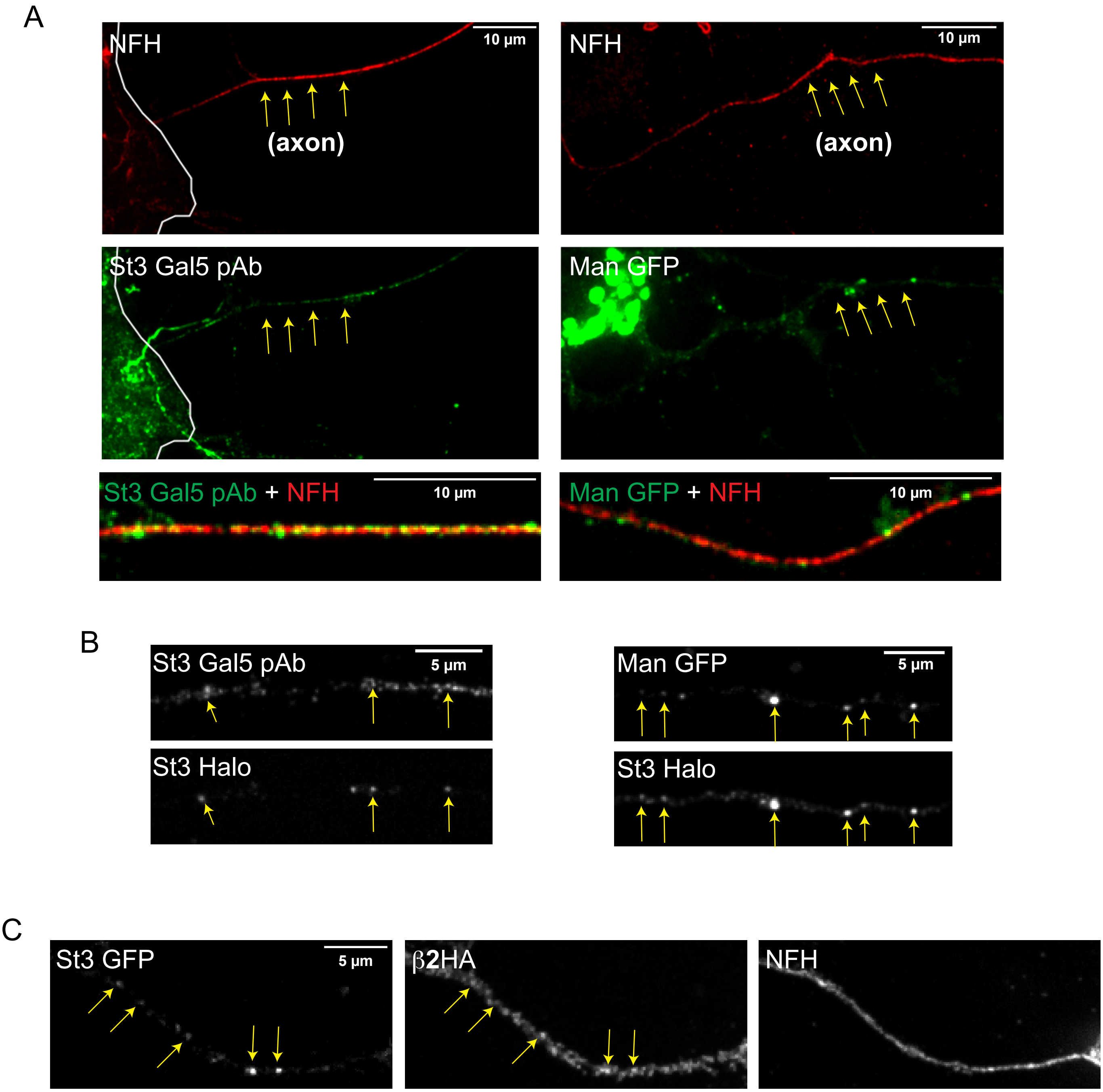
A
NFH
10 μm
(axon)
NFH
10 μm
(axon)
St3 Gal5 pAb
Man GFP
St3 Gal5 pAb + NFH
10 μm
Man GFP + NFH
10 μm
B
St3 Gal5 pAb
5 μm
St3 Halo
Man GFP
5 μm
St3 Halo
C
St3 GFP
5 μm
β2HA
NFH

A

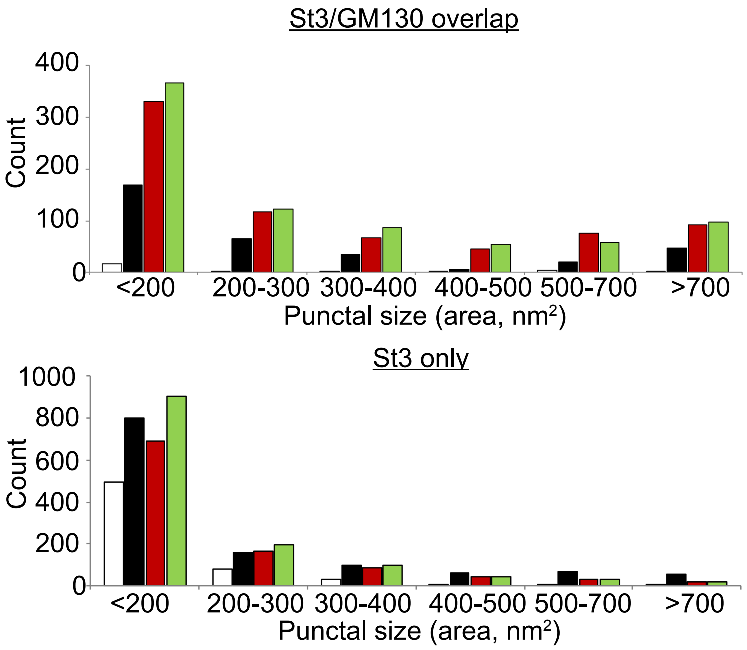
St3/GM130 overlap
Count
400
300
200
100
0
<200
200-300
300-400
400-500
500-700
>700
Punctal size (area, nm²)
St3 only
1000
800
600
400
200
0
Count
<200
200-300
300-400
400-500
500-700
>700
Punctal size (area, nm²)

B

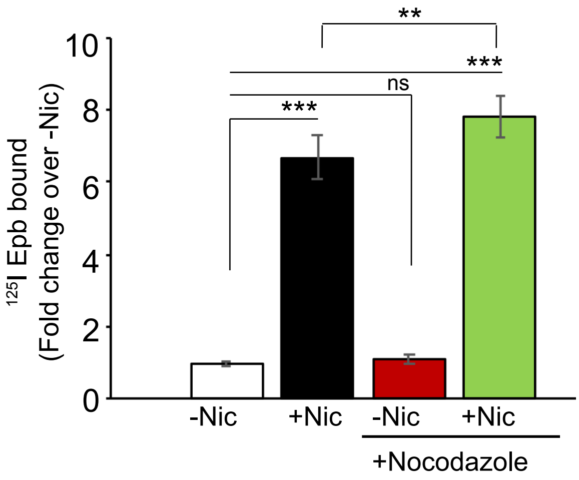
**
***
ns
***
10
8
6
4
2
0
125I Epb bound
(Fold change over -Nic)
-Nic
+Nic
-Nic
+Nic
+Nocodazole

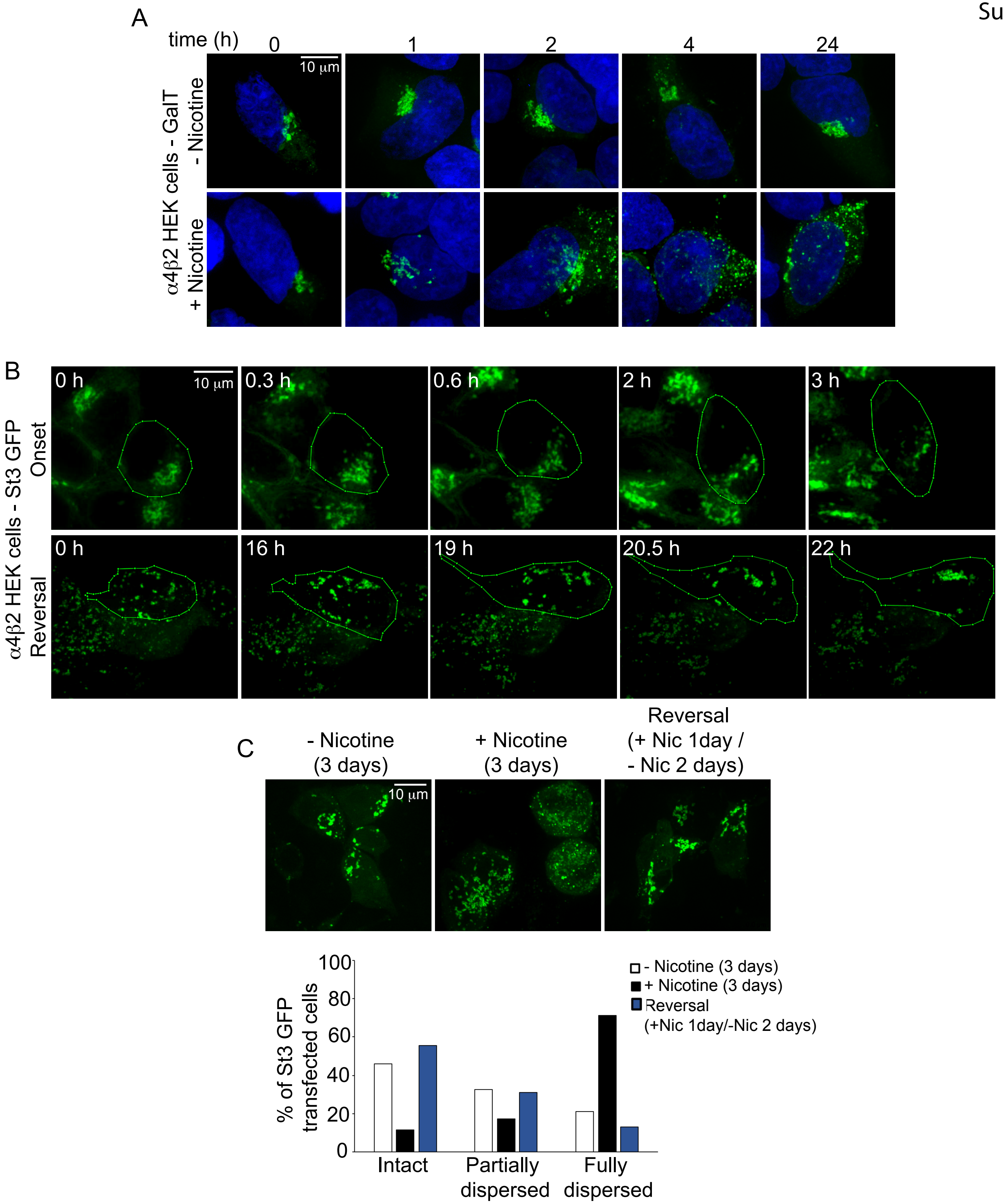
A
time (h)
0
1
2
4
24
10 μm
α4β2 HEK cells - GalT
- Nicotine
+ Nicotine
B
α4β2 HEK cells - St3 GFP
Onset
Reversal
0 h
10 μm
0.3 h
0.6 h
2 h
3 h
0 h
16 h
19 h
20.5 h
22 h
C
- Nicotine (3 days)
+ Nicotine (3 days)
Reversal (+ Nic 1day / - Nic 2 days)
10 μm
% of St3 GFP transfected cells
100
80
60
40
20
0
Intact
Partially dispersed
Fully dispersed
- Nicotine (3 days)
+ Nicotine (3 days)
Reversal (+Nic 1day/-Nic 2 days)

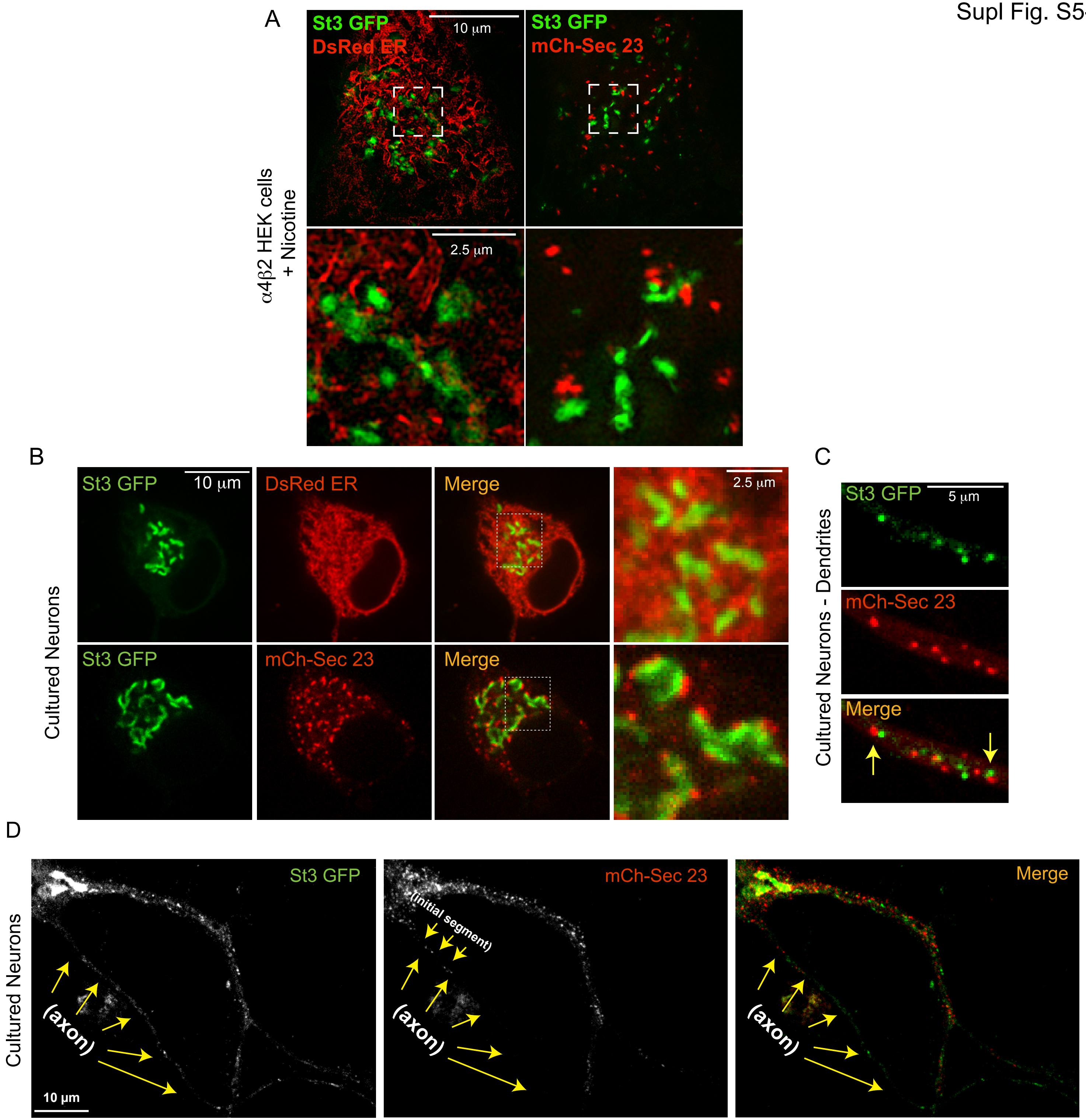
A
St3 GFP
DsRed ER
10 μm
St3 GFP
mCh-Sec 23
α4β2 HEK cells
+ Nicotine
2.5 μm
B
Cultured Neurons
St3 GFP
10 μm
DsRed ER
Merge
2.5 μm
St3 GFP
mCh-Sec 23
Merge
C
Cultured Neurons - Dendrites
St3 GFP
5 μm
mCh-Sec 23
Merge
D
Cultured Neurons
St3 GFP
(axon)
10 μm
mCh-Sec 23
(initial segment)
(axon)
Merge
(axon)

A

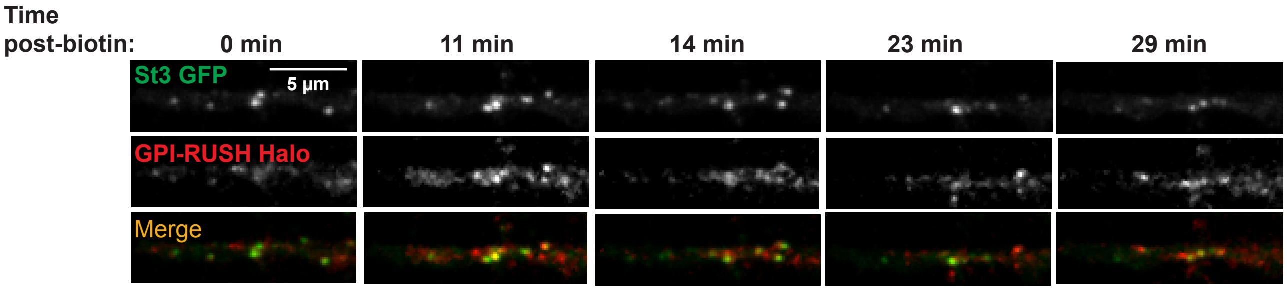
Time
post-biotin:
0 min
11 min
14 min
23 min
29 min
St3 GFP
5 µm
GPI-RUSH Halo
Merge
